# A multimodal foundation model linking histopathology and DNA methylation

**DOI:** 10.64898/2026.06.11.731518

**Authors:** Dannong Wang, Jessica Zhang, Chen Chen, Wei Zhang, Song Wang, Yanda Meng, Guru Sonpavde, Craig Horbinski, Yu Tian

## Abstract

Hematoxylin and eosin (H&E) slides are routinely available in cancer care, but molecular profiling often requires additional tissue processing and turnaround time. We introduce HistoMethyl, a DNA methylation-aware pathology foundation model that aligns whole-slide histopathology with matched genome-scale methylation beta-value profiles during pre-training while requiring only an H&E slide at inference. We evaluated HistoMethyl across four task groups: gene mutation prediction, morphology-associated classification, overall survival prediction, and direct DNA methylation beta-value recovery. Evaluation spanned TCGA cross-validation, a disease-held-out lower-grade glioma cohort, and external cohort validation in CPTAC glioblastoma and SurGen rectal adenocarcinoma, totaling 14 cancer cohorts and 81 gene mutation tasks. The best-performing configuration improved mean mutation AUROC by 5.03 percentage points. By converting DNA methylation supervision into an H&E-only representation, HistoMethyl could support an early molecular triage layer that helps prioritize cases for confirmatory sequencing, methylation profiling, immunohistochemistry, and molecular tumor board review. These image-only predictions are intended to accelerate downstream molecular testing and tissue allocation while leaving final diagnosis and treatment selection anchored in validated molecular assays.

## 1 Introduction

Routine hematoxylin and eosin (H&E) histopathology remains the most widely available tissue-based readout for cancer diagnosis. Recent whole-slide foundation models have shown that large-scale pre-training can produce transferable histopathology representations across diagnostic, prognostic, and molecular prediction tasks [1–3]. In parallel, multimodal pathology models have demonstrated that histology can be aligned with molecular measurements such as transcriptomics and spatial transcriptomics, enabling prediction of gene expression, spatial tissue organization, mutations, and clinical outcomes from morphology [4–6]. However, most existing approaches either learn image-only visual representations, train task-specific molecular predictors, or use transcriptomic supervision. DNA methylation has received comparatively less attention as a supervision signal for reusable whole-slide representations, despite its central role in defining tumor lineage, cell state, and clinically meaningful epigenetic subtypes [4–8].

DNA methylation is particularly relevant for molecularly informed pathology because it captures stable epigenomic programs linked to tumor lineage, cell state, and disease classification. Across cancer types, methylation patterns have been associated with tumor cell-of-origin, molecular subtype, prognosis, and therapeutic vulnerabilities, supporting DNA methylation as a biologically structured signal for pan-cancer representation learning [9]. Its clinical value is especially well established in central nervous system tumors, where methylation-based classification has become an important component of integrated diagnosis and diagnostic refinement [10–12]. Yet methylation profiling is not universally available, may require referral to specialized laboratories, and can delay clinical interpretation in settings where rapid molecular information is needed [13]. These constraints motivate a complementary strategy: rather than replacing methylation profiling, DNA methylation profiles can be used as biologically structured supervision for learning H&E slide representations that capture methylation-associated morphology.

Here, we introduce HistoMethyl, a DNA methylation-enhanced pathology foundation model that learns aligned representations of histopathology whole-slide images and matched DNA methylation profiles. HistoMethyl uses paired whole-slide images and methylation beta-value profiles from 7,769 unique patients to learn a shared image-methylation representation space. At inference, HistoMethyl requires only an H&E whole-slide image. This design allows HistoMethyl to recover methylation-associated information from morphology while still preserving the visual signal needed for routine pathology analysis. Rather than targeting a single biomarker or cancer type, HistoMethyl is intended as a general-purpose methylation-aware whole-slide foundation model whose image representations are explicitly shaped by epigenomic tumor state.

To test whether our methylation-enhanced pathology foundation model improves routine H&E representations, we evaluated HistoMethyl across mutation prediction, morphology-associated classification, overall survival prediction, and DNA methylation beta-value recovery. Evaluation spanned cross-validation within The Cancer Genome Atlas Program (TCGA) [14], a full disease-held-out lower-grade glioma (LGG) cohort from TCGA, and external cohort validation in CPTAC-GBM (glioblastoma) [15] and rectal adenocarcinoma (READ) from the SurGen SR386 dataset [16], totaling 14 cancer cohorts and 81 unique gene mutation prediction tasks. Across these settings, HistoMethyl improved aggregate performance over image-only pathology foundation models: the best-performing configuration improved mean mutation AUROC by 5.0 percentage points, morphology-associated AUROC by 1.2 percentage points, survival C-index by 0.023, and flattened DNAm Pearson correlation by 0.046. These findings support DNA methylation as an epigenomic supervision signal for learning molecularly informative H&E slide representations and position HistoMethyl as a representation-learning and decision-support framework rather than a replacement for clinical molecular testing. More broadly, HistoMethyl points toward a future of computational pathology in which molecular assays are not only endpoints to be predicted from images, but also biologically structured supervision for learning pathology representations that support precision oncology while retaining practical image-only inference.

## 2 Results

### 2.1 A DNA methylation-aware whole-slide representation model for image-only pathology inference

HistoMethyl uses matched DNA methylation profiles as molecular supervision during pre-training to learn image-only whole-slide representations for downstream clinical and genomic prediction. The model consists of two aligned branches: an image branch that encodes hematoxylin and eosin whole-slide images and a methylation branch that encodes matched DNA methylation beta-value profiles. We instantiated the image branch using two pathology foundation model backbones, Prov-GigaPath [3] and Feather-24k [17], and used CpGPT [18], a pre-trained methylation encoder, to represent CpG identities and beta values. The two branches were projected into a shared representation space and trained using a contrastive image-methylation alignment objective on paired cases.

#### Data

HistoMethyl was pre-trained once per image backbone using a downstream-task-agnostic image-DNAm alignment objective on paired H&E whole-slide images and DNA methylation profiles from TCGA [14], with the entire LGG cohort excluded from pre-training. After pre-training, the methylation branch was removed, and all downstream models used H&E-derived slide representations only. We evaluated the pre-trained image encoder in three complementary settings: TCGA in-corpus representation evaluation, disease-held-out evaluation on the LGG cohort (364 patients), and external cohort evaluation in CPTAC-GBM [15] (99 patients) and SurGen READ [16] (145 patients). In all settings, task-specific classification/regression heads were trained and evaluated using patient-level stratified five-fold cross-validation across three independent runs. The foundation encoder itself was trained once, reflecting its intended use as a reusable representation model rather than a split-specific supervised model. Thus, the TCGA in-corpus setting evaluates in-sample downstream label generalization, whereas TCGA-LGG, CPTAC-GBM, and SurGen READ assess whether the same pre-trained representation remains useful beyond the pre-training disease cohort or data source. This design evaluates whether methylation-aware pre-training improves image-only representations across internal, disease-held-out, and external evaluation settings.

#### Pre-training analysis

Figure 1b,c shows the pre-training pipeline, in which slide embeddings and DNAm embeddings are projected by multilayer perceptrons and aligned using a contrastive loss. We first evaluated whether this task-agnostic alignment improved the organization of the slide representation before any downstream supervision was introduced. Figure 2a shows t-SNE visualizations of the base and methylation-aligned slide embeddings, colored by the 12 largest cancer cohorts. Compared with the baseline embeddings, the methylation-aligned representations showed clearer cohort-level structure, consistent with a methylation-associated reorganization of cohort-level slide embeddings. Figure 2b further quantifies image-DNAm alignment using retrieval recall. For the baseline models, each slide embedding from the original image backbone was used to retrieve its matched DNAm embedding from the validation set. For HistoMethyl, the methylation-aligned slide embedding was used for the same retrieval task. Recall@k reports the fraction of slides for which the matched DNAm profile was ranked among the top k retrieved methylation embeddings, with higher recall indicating stronger cross-modal alignment. Averaged across the four evaluated cohorts, HistoMethyl improved Recall@1, Recall@5, and Recall@10 for both backbones. Together, these analyses show that methylation-aware pre-training improves cross-modal correspondence before downstream task adaptation, supporting the use of DNAm as a molecular supervision signal for slide representation learning.

**Fig. 1.**
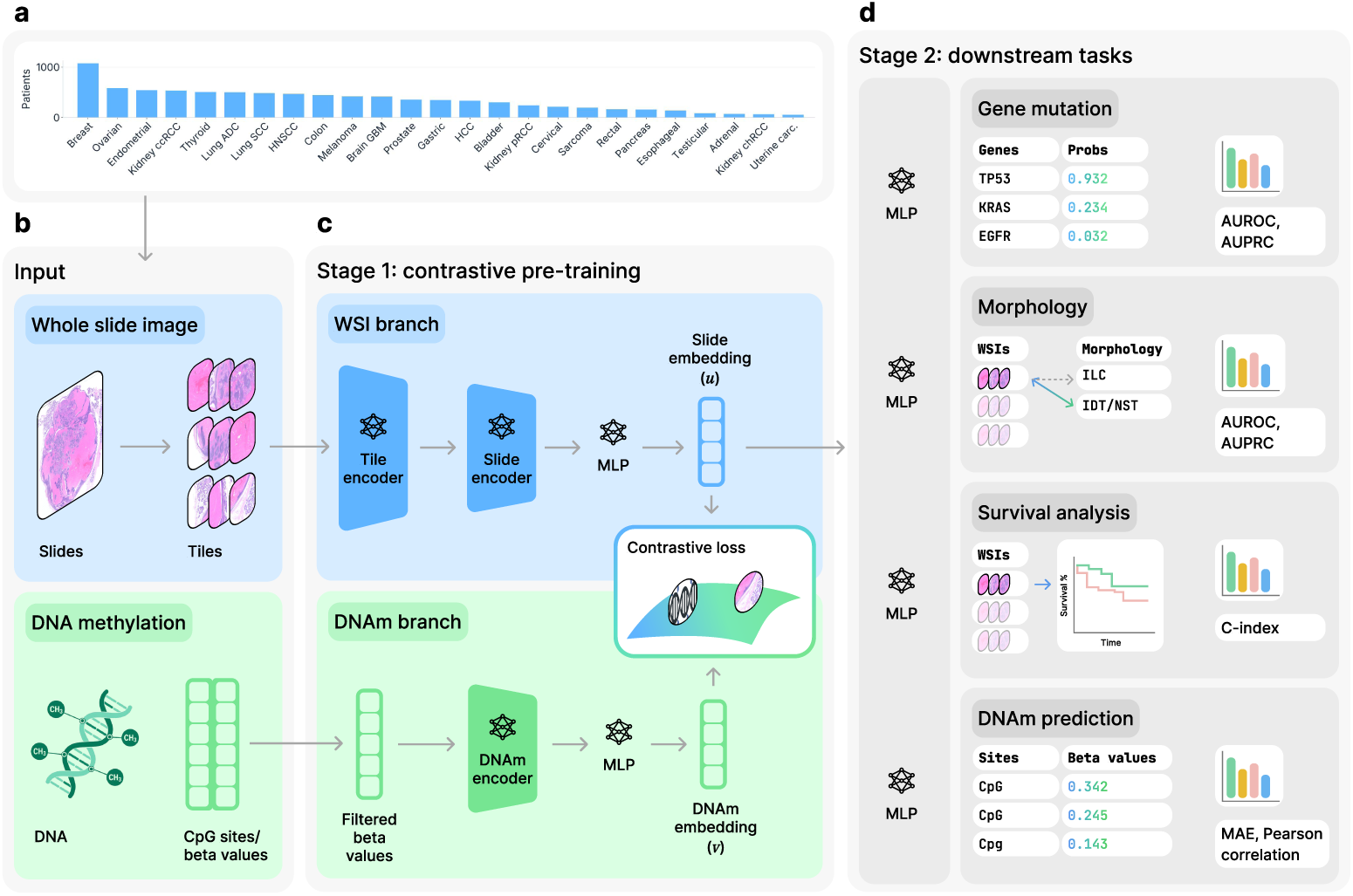
HistoMethyl overview. **a**, Cancer cohorts from TCGA [14] used for HistoMethyl pre-training. **b**, Pre-training inputs, including WSIs divided into tiles and paired DNAm beta values corresponding to each case. **c**, Pre-training with contrastive alignment. WSI and DNAm embeddings are extracted by their respective encoders, projected by MLPs, and aligned with a contrastive loss. **d**, Downstream task adaptation. HistoMethyl uses task-specific supervised models to adapt aligned slide embeddings to downstream tasks, including gene mutation, morphology, survival analysis, and DNAm prediction.

**Fig. 2.**
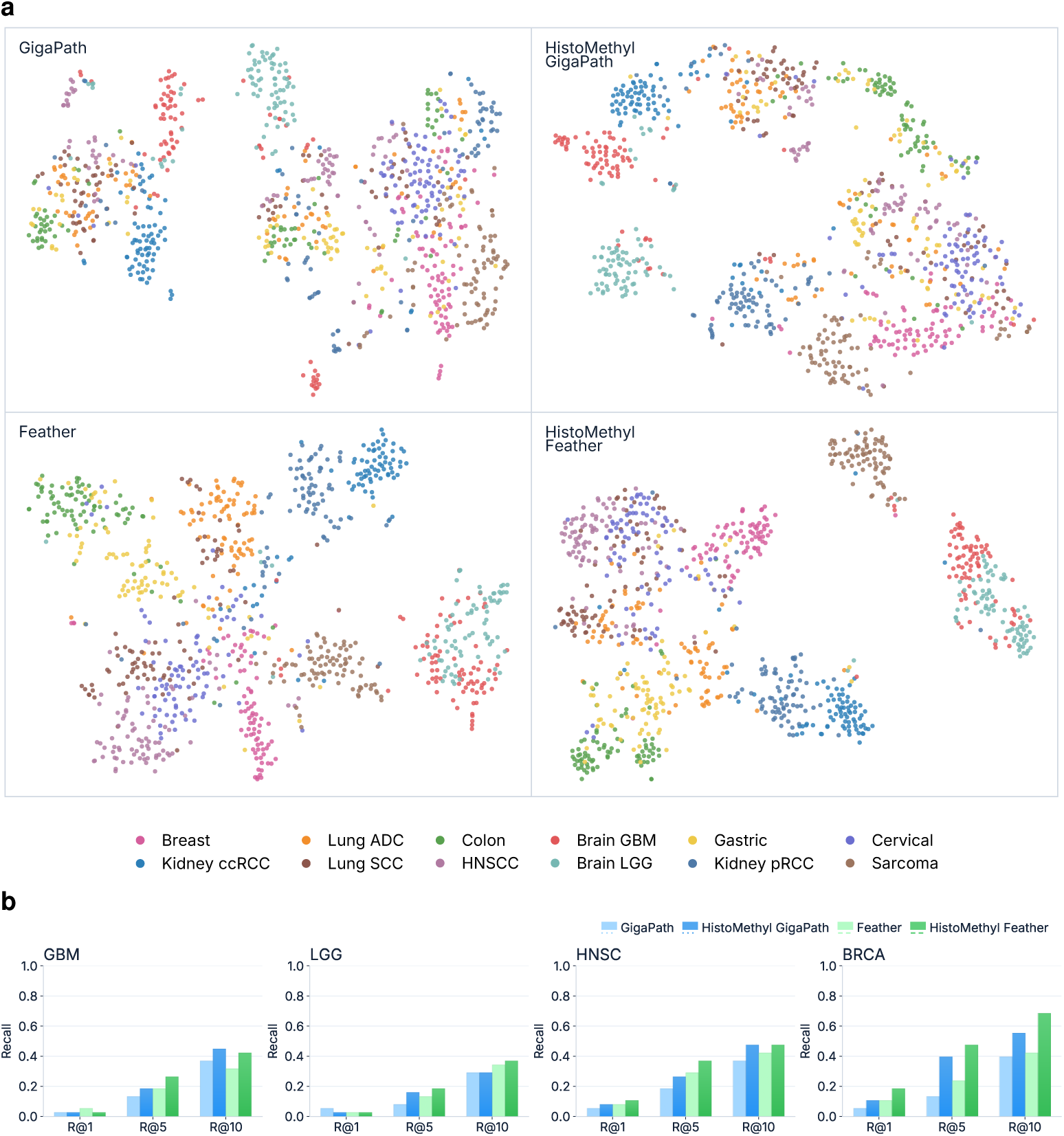
Representation-level assessment of methylation alignment. **a**, t-SNE visualization of slide embeddings from the four slide embedding variants, colored by cancer type. **b**, Image-to-DNAm retrieval across GBM, LGG, HNSC, and BRCA using matched slide and methylation embeddings, shown as Recall@1, Recall@5, and Recall@10. These analyses use pre-trained embeddings before supervised downstream task adaptation, providing a direct assessment of whether image-methylation contrastive alignment changed the representation space.

#### Downstream task adaptation

After pre-training, only the image branch was retained for evaluation, so all downstream tasks used H&E-derived slide embeddings alone without requiring methylation profiles at inference. We evaluated HistoMethyl across four task families chosen to test complementary aspects of a methylation-aware pathology representation. Mutation prediction and DNAm beta-value prediction directly evaluate whether methylation alignment enriches the image embedding for molecular and epigenomic information closely related to the pre-training signal, as DNA methylation captures tumor lineage, molecular subtype, and cancer-associated regulatory programs across tumor types [9, 10, 19]. Morphology-associated prediction and overall survival analysis evaluate whether this alignment preserves broader whole-slide information needed for conventional pathology interpretation and clinically relevant prognostic modeling [8, 20, 21]. For classification tasks, supervised heads were trained with patient-level stratified five-fold cross-validation repeated across three independent runs, while the pre-trained foundation encoder remained fixed. Survival analysis used Cox models on fixed embeddings, whereas DNAm beta-value prediction used image-only regression with slide-encoder fine-tuning. Together, these four task families provide a comprehensive evaluation of whether HistoMethyl improves molecularly aligned representations without sacrificing general histopathologic and prognostic utility. Full results are provided in Supplementary Section 1.

### 2.2 Methylation-aware pre-training improves mutation prediction across cohorts

HistoMethyl produced its strongest and broadest gains in gene mutation prediction, improving image-only mutation classification across TCGA [14] cross-validation cohorts, the fully held-out LGG cohort, and external diagnostic-slide evaluation cohorts. We evaluated whether methylation-aware pre-training improved the ability of H&E-derived slide embeddings to predict somatic mutation status. In Figure 3, mutation performance was summarized as mean AUROC within each cohort, with each HistoMethyl model compared against its matched image-only backbone.

**Fig. 3.**
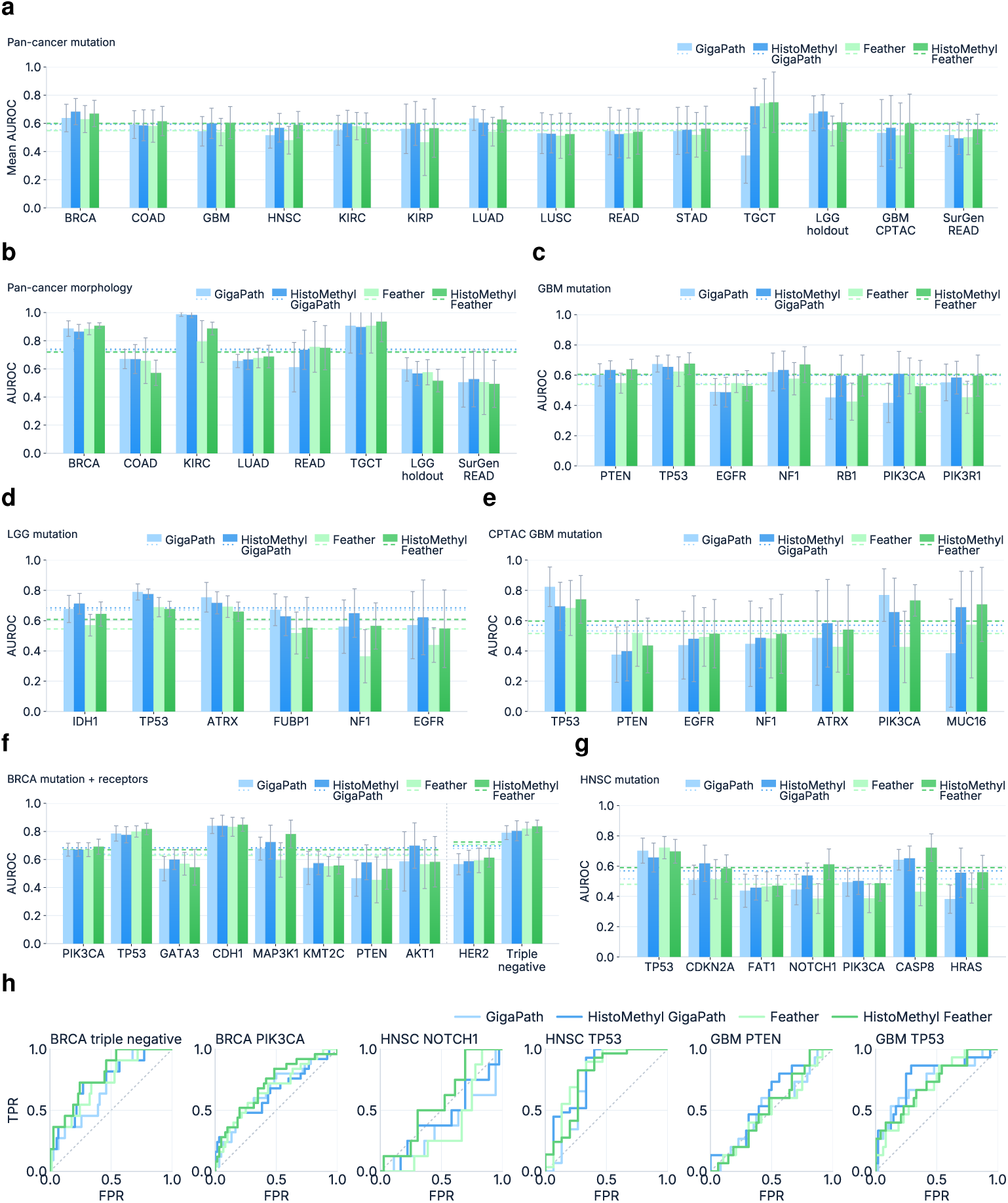
Gene mutation prediction and morphology tasks. **a**, Mean AUROC across cohort-specific mutation prediction tasks. **b**, Morphology prediction AUROC across cohorts. **c–e**, Detailed mutation prediction results for GBM, lower-grade glioma (LGG), and CPTAC GBM. **f**, Breast cancer (BRCA) mutation and receptor prediction, shown with HER2 and triple-negative receptor endpoints as a representative non-CNS cohort example. **g**, Head and neck squamous cell carcinoma (HNSC) mutation prediction, shown as a second representative non-CNS cohort example. **h**, Representative held-out ROC curves for selected BRCA, HNSC, and GBM tasks. Error bars indicate standard deviation across held-out cross-validation folds and repeated runs where applicable. Horizontal dashed lines indicate within-panel model means, and in **f** the mean lines are computed separately for the mutation and receptor groups.

Across 81 gene mutation prediction tasks in 14 cancer cohorts, HistoMethyl improved mutation prediction for both backbone families. For this aggregate analysis, AUROCs were first averaged across mutation genes within each cohort, and cohort means were then averaged equally across cohorts. HistoMethyl GigaPath improved mean mutation AUROC by 4.08 percentage points over GigaPath [3], while HistoMethyl Feather improved mean mutation AUROC by 5.03 percentage points over Feather [17]. A HistoMethyl model achieved the highest cohort-level mean mutation AUROC in 11 of 14 cohorts. Using a two-sided paired Wilcoxon signed-rank test across cohort-level mean mutation AUROC values, the improvement was statistically significant for both matched backbones, with HistoMethyl GigaPath outperforming GigaPath at *p <* 0.05 and HistoMethyl Feather outperforming Feather at *p <* 0.001.

The detailed disease panels further show that these gains were not driven by a single cancer type or isolated gene. In GBM, HistoMethyl improved mean gene-level AUROC by 5.61 percentage points for the GigaPath backbone and 6.68 percentage points for the Feather backbone. Shared gains were observed for NF1, PTEN, RB1, and PIK3R1, with additional improvement for PIK3CA in the GigaPath-based model and TP53 in the Feather-based model. In the fully held-out LGG cohort, which was excluded from methylation-alignment pre-training, HistoMethyl GigaPath improved mean mutation AUROC by 1.37 percentage points and HistoMethyl Feather improved mean mutation AUROC by 6.20 percentage points. Both HistoMethyl variants improved prediction of IDH1, NF1, and EGFR, and HistoMethyl Feather additionally improved FUBP1 prediction, supporting transfer to a disease cohort not used during contrastive pre-training.

HistoMethyl also improved mutation prediction in clinically relevant non-CNS cohorts. In BRCA, HistoMethyl GigaPath improved mean mutation AUROC by 4.58 percentage points and HistoMethyl Feather improved mean mutation AUROC by 3.98 percentage points, with shared gains for AKT1, CDH1, KMT2C, MAP3K1, PIK3CA, and PTEN. In HNSC, both HistoMethyl variants improved six of seven displayed mutation endpoints, including CASP8, CDKN2A, FAT1, HRAS, NOTCH1, and PIK3CA, with especially strong mean gains for HistoMethyl Feather.

External-cohort validation provided an important test of whether methylation-aware representations generalized beyond TCGA. In CPTAC-GBM, HistoMethyl improved mean mutation AUROC for both backbones, with HistoMethyl Feather improving mean AUROC by 8.28 percentage points. Both HistoMethyl variants improved ATRX, EGFR, MUC16, and NF1 prediction in this cohort. In the SurGen READ cohort, HistoMethyl Feather improved RAS mutation prediction by 5.76 percentage points over Feather, providing additional evidence of external diagnostic-slide transfer.

Together, these results indicate that methylation-aware pre-training improves mutation prediction at both the cohort and gene level, with the strongest evidence observed in molecularly relevant endpoints and retained in external-cohort validation settings.

### 2.3 Methylation alignment extends to morphology and survival prediction

Beyond mutation prediction, HistoMethyl preserved strong morphology-associated performance and improved aggregate prognostic prediction, indicating that methylation alignment enhanced molecular informativeness without sacrificing broader slide-level utility. Morphology-associated predictions test whether alignment to DNAm retained visually grounded histologic information. As shown in Figure 3b, HistoMethyl GigaPath improved mean morphology AUROC by 1.20 percentage points across the eight cohorts, while HistoMethyl Feather maintained the strong performance of the baseline models. These results indicate that methylation-aware pre-training can strengthen histologic phenotype prediction in selected disease settings while maintaining competitive performance across diverse morphology tasks.

Figure 4a details overall survival prediction using image-only slide embeddings. HistoMethyl Feather improved mean C-index by 0.023 across the 11 displayed cohorts, with cohort-level gains in BRCA, GBM, HNSC, KIRC, and READ. HistoMethyl GigaPath also improved survival prediction in GBM, LUAD, and READ. Because overall survival reflects many clinical, treatment, and non-image factors, these gains suggest that methylation-aware pre-training can capture prognostically relevant morphology when such signal is present in the slide representation.

**Fig. 4.**
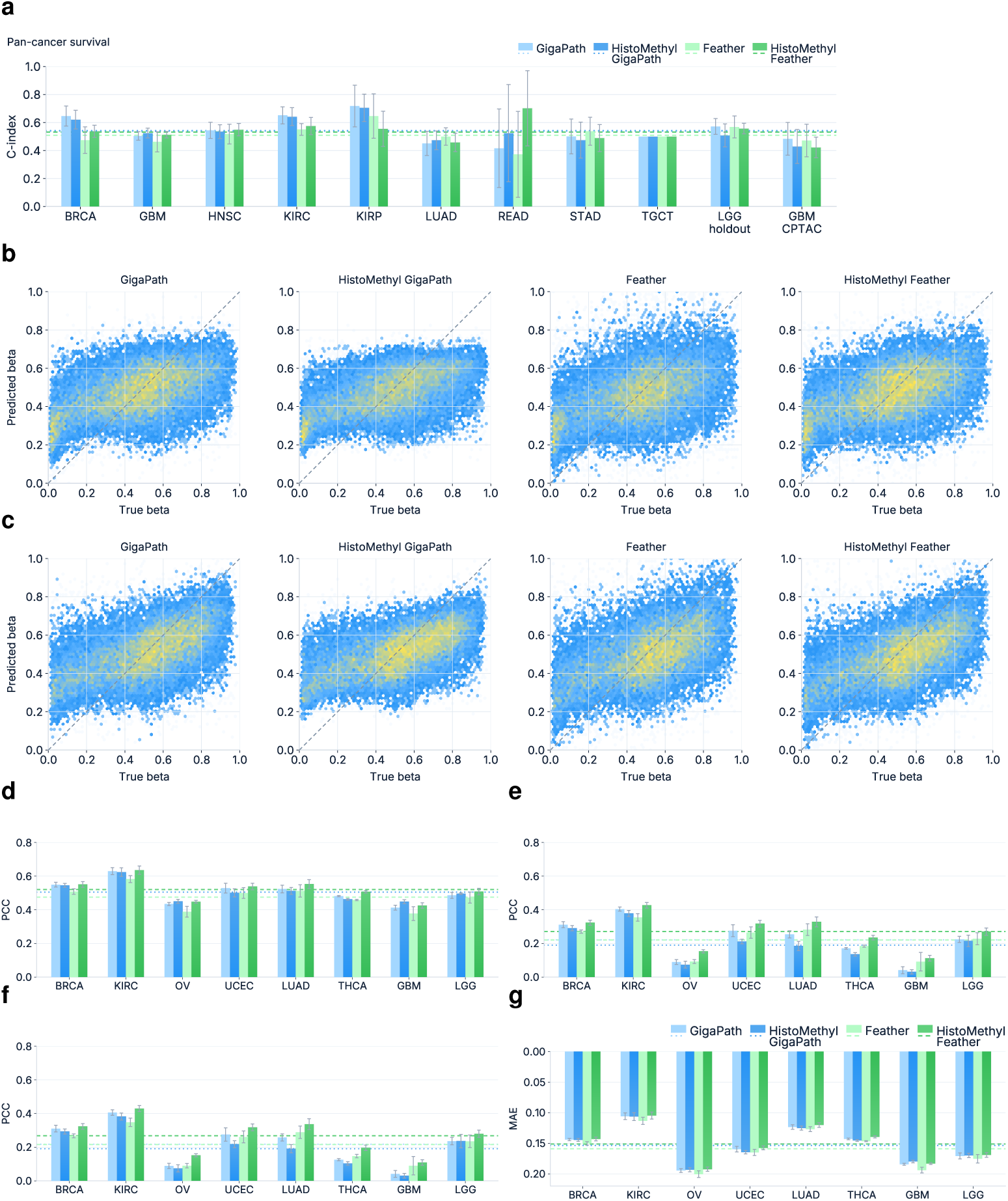
Survival analysis and DNA methylation beta-value prediction. **a**, Mean concordance index for overall survival prediction. **b,c**, Predicted versus measured DNA methylation beta values for GBM and LGG. **d**, Flattened Pearson correlation coefficient across all eight evaluated DNAm prediction cohorts. **e**, Mean site-wise Pearson correlation coefficient. **f**, Median site-wise Pearson correlation coefficient. **g**, Mean absolute error, plotted with an inverted y-axis so lower error appears higher. Error bars indicate cross-validation standard deviation, and horizontal dashed lines indicate mean model performance across the displayed cohorts.

Taken together, the morphology and survival analyses show that the benefits of HistoMethyl are not limited to direct genomic classification. Methylation alignment preserves and even improves on broader histologic and prognostic tasks, supporting the use of DNAm as a general molecular supervision signal for image-only pathology representation learning.

### 2.4 HistoMethyl enriches image embeddings for DNAm beta-value prediction

Direct DNAm beta-value prediction provided a direct test of whether methylation-aware pre-training enriched image-only slide embeddings with recoverable epigenomic signal. We evaluated this task across eight cohorts with available DNAm prediction results. Figure 4 shows representative scatter plots for GBM and LGG and summarizes all eight cohorts using flattened Pearson correlation, mean and median site-wise Pearson correlation, and mean absolute error, capturing global beta-value recovery, CpG-level recovery, and calibration error.

Across all eight evaluated cohorts, HistoMethyl Feather showed the clearest and most consistent improvement over the matched Feather baseline. Relative to Feather, HistoMethyl Feather improved flattened Pearson correlation by 4.6 percentage points, mean site-wise Pearson correlation by 5.0 percentage points, median site-wise Pearson correlation by 5.2 percentage points, and reduced mean absolute error by 0.008 on average, with particularly strong site-wise correlation improvements in KIRC and OV, while GBM and LGG showed improved global correlation and lower prediction error despite more modest absolute site-wise correlations. Across the eight evaluated cohorts, these improvements were statistically significant for all reported metrics using a two-sided paired Wilcoxon signed-rank test, with *p <* 0.01.

The HistoMethyl model using the GigaPath backbone [3] also showed cohort-level gains in direct beta-value prediction, with improvements in flattened correlation and mean absolute error in GBM, LGG, and OV. These results indicate that methylation alignment can enrich image-only embeddings with DNAm-associated morphology, with the strongest aggregate improvement observed for the Feather backbone.

### 2.5 Occlusion sensitivity analyses provide qualitative examples of mutation prediction

Occlusion-based heatmaps [22] were used to compare the tissue regions supporting each mutation classifier. This perturbation strategy, developed to measure prediction changes after masking local image regions and later used in medical AI studies across pathology [23–25], mammography [26], and neuroimaging [27], was selected because it provides task-specific evidence rather than encoder-level attention. This distinction is important in our setting, since classification probes operate on frozen slide embeddings and attention- or CLAM-style maps [28] may reflect shared tissue selection or general slide salience rather than the mutation-specific decision boundary. For each selected slide, local tissue patch groups were masked, the slide embedding and mutation score were recomputed, and regional importance was assigned according to the reduction in positive-class score. The resulting maps therefore highlight tissue regions whose removal most weakened a specific mutation prediction, enabling qualitative comparison of whether HistoMethyl and image-only baselines relied on coherent and concordant spatial evidence.

We selected representative examples in Figure 5 spanning CNS tumors, head and neck cancer, and external CPTAC GBM to show whether the observed behavior was limited to a single disease setting or appeared across multiple evaluation contexts. In these examples, methylation-aligned models often focus on overlapping tissue regions, while baseline encoders are more diffuse or emphasize different areas of the same slide. This analysis is qualitative and should not be interpreted as pathologist-validated localization of molecular clones; instead, it provides supporting evidence that methylation-aligned pre-training can alter the spatial evidence captured by the slide representation.

**Fig. 5.**
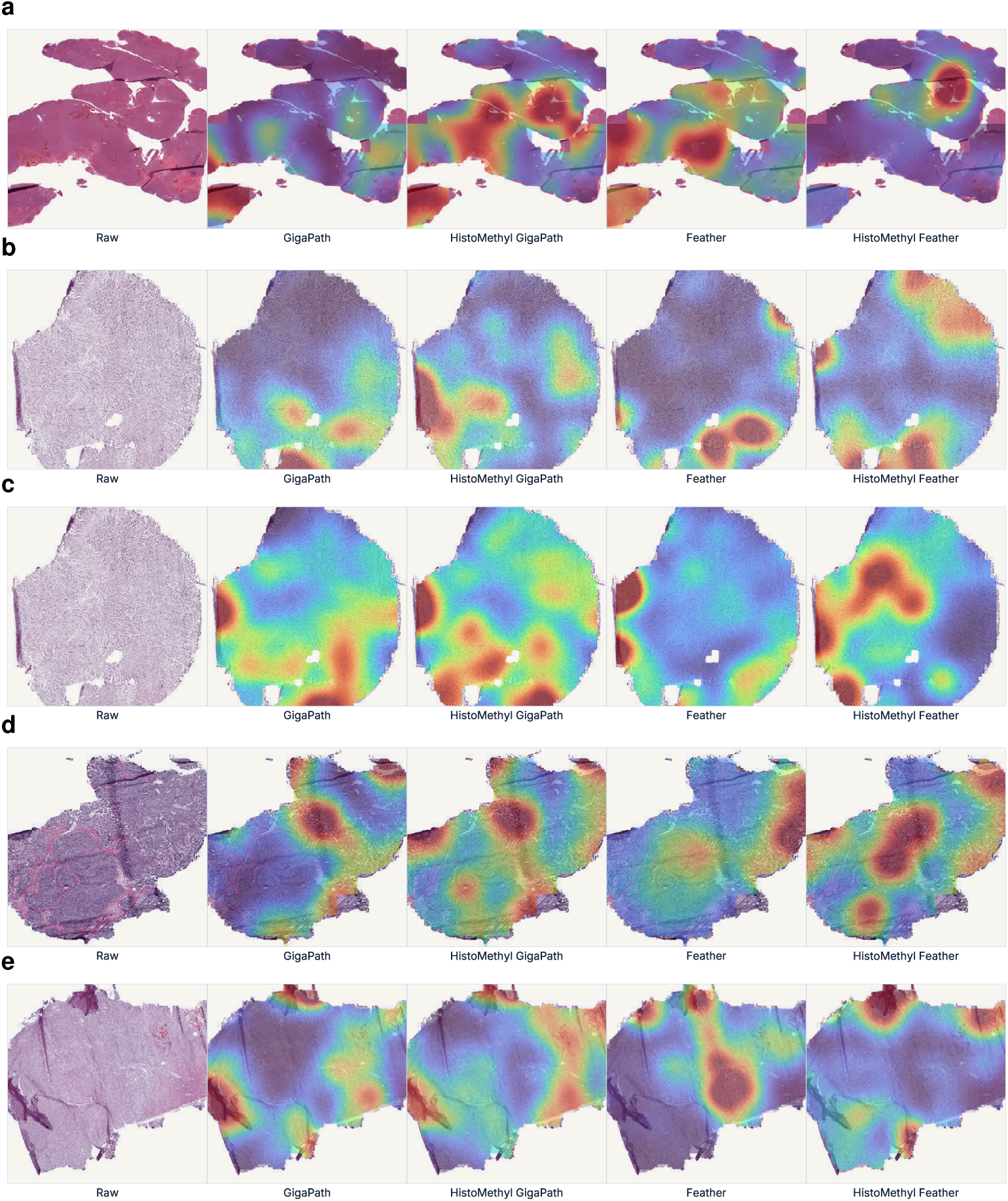
Representative occlusion-based mutation heatmaps. Each row shows the raw H&E thumbnail followed by heatmaps from GigaPath [3], HistoMethyl GigaPath, Feather [17], and HistoMethyl Feather. **a**, CPTAC GBM NF1. **b**, GBM EGFR. **c**, GBM TP53. **d**, HNSC CDKN2A. **e**, LGG TP53. Higher-intensity regions indicate tissue areas whose masking reduced the mutation probe decision score most strongly.

## 3 Discussion

HistoMethyl demonstrates that DNA methylation can be used not only as a molecular endpoint, but also as a biologically structured supervision signal for learning clinically useful whole-slide representations. Across downstream evaluations, methylation alignment improved image-only embeddings across mutation prediction, morphology-associated classification, survival analysis, and direct DNAm beta-value recovery, with the strongest and broadest gains in gene mutation prediction. This pattern is clinically meaningful because the model is not simply reconstructing the pre-training modality; rather, DNAm supervision enriches the H&E representation for molecular phenotypes that are relevant to diagnostic classification and prognostic assessment [9, 19]. The pan-cancer evaluation is therefore essential to the interpretation of HistoMethyl. In CNS tumors, methylation profiling has already shown direct diagnostic utility by refining histopathologic classification and supporting integrated molecular diagnosis [10]. HistoMethyl extends this clinical paradigm beyond CNS by showing that methylation-aware supervision can improve image-only representations across multiple cancer cohorts, rather than serving only as a disease-specific neuro-oncology predictor. Thus, CNS disease provides a clinically mature example of methylation-guided diagnosis, while the broader contribution of HistoMethyl is a pan-cancer framework in which epigenomic tumor state shapes H&E slide representations.

The most immediate clinical implication is earlier molecular triage from an H&E slide before confirmatory molecular results are available. In current oncology workflows, molecular testing requires specimen review, tumor selection, tissue handling, nucleic-acid extraction, assay processing, interpretation, and reporting, all of which can delay molecularly guided care decisions [29]. Because molecular findings guide diagnosis, targeted therapy selection, clinical-trial eligibility, and integrated tumor classification, earlier identification of cases likely to carry clinically relevant molecular states could help prioritize confirmatory testing rather than replace it [30]. In this context, HistoMethyl could serve as a rapid and practical decision-support tool: cases with high predicted probability of a clinically relevant molecular alteration or epigenomic state could be prioritized for expedited sequencing, methylation profiling, immunohistochemistry, and molecular tumor board discussion when genomic findings require multidisciplinary interpretation [31, 32]. The morphology and survival results further indicate that this molecular enrichment does not eliminate conventional slide-level information, which is important because a clinically useful pathology representation must preserve diagnostic tissue context while adding molecular signal.

The frozen-slide design is central to this workflow interpretation. HistoMethyl aligns frozen H&E slides with DNAm profiles because TCGA [14] genomic and epigenomic data were generated from frozen tissue, and TCGA frozen-section images are linked to tissue submitted for molecular characterization [33, 34]. This pairing makes the image-methylation objective more biologically grounded than aligning a molecular profile from one tissue region with a diagnostic slide from another block. It also matches the clinical motivation: frozen sections can be generated during intraoperative or rapid-consultation workflows, whereas routine diagnostic processing and molecular testing commonly add additional time [29, 31, 35]. A DNAm-informed embedding derived directly from a frozen slide could therefore provide an early molecularly enriched representation near the point of initial pathology review. Conversely, external-cohort validation on CPTAC glioblastoma [15] and SurGen colorectal [16] diagnostic slides tests the complementary deployment question: whether a representation learned from frozen-slide-to-DNAm alignment can transfer to routine diagnostic-slide settings despite differences in tissue processing, artifacts, staining, scanning, and sampling. The external results therefore strengthen the clinical interpretation by linking a rapid frozen-section motivation with broader diagnostic-slide applicability.

In conclusion, HistoMethyl supports a model of computational pathology in which molecular assays are used to train more informative slide representations, while inference remains practical and image-only. Its contribution is not simply the improved performance over image-only baselines, but a route for bringing epigenomic context closer to the earliest tissue-based decision point in cancer care. In prospective use, such a system could serve as an early, tissue-preserving triage layer that helps prioritize confirmatory molecular testing, guide tissue allocation, and coordinate downstream clinical review while leaving final diagnosis and treatment selection anchored in validated assays.

## 4 Methods

### 4.1 Data

HistoMethyl was developed in two stages: contrastive pre-training on paired histopathology whole-slide images and DNA methylation profiles, followed by image-only downstream evaluation. For pre-training, we used pan-cancer cohorts from The Cancer Genome Atlas (TCGA) [14], which provides large matched molecular and clinical datasets across many tumor types and is widely used for pan-cancer computational pathology studies. To evaluate the generalizability of our model specifically in a CNS disease setting, the lower-grade glioma (LGG) cohort was excluded entirely from contrastive pre-training so that it could serve as a held-out disease cohort in downstream evaluation. Similarly, the CPTAC glioblastoma (GBM) cohort was not used during pre-training and was reserved as an external evaluation cohort. This design separated in-distribution transfer from disease-held-out and external generalization.

The central clinical motivation of the study was neuro-oncology. We therefore placed particular emphasis on GBM, LGG, and CPTAC GBM, because DNA methylation already plays an established role in CNS tumor classification and diagnostic refinement [10, 36, 37]. To show that gains were not restricted to brain tumors, we selected additional cohorts spanning several disease groups: gastrointestinal adenocarcinomas, including colon adenocarcinoma (COAD), rectum adenocarcinoma (TCGA-READ and SurGen-READ), and stomach adenocarcinoma (STAD); thoracic tumors, including lung adenocarcinoma (LUAD) and lung squamous cell carcinoma (LUSC); breast cancer (BRCA); head and neck squamous cell carcinoma (HNSC); renal tumors, including kidney renal clear cell carcinoma (KIRC) and kidney renal papillary cell carcinoma (KIRP); and testicular germ cell tumor (TGCT). Together, they cover clinically distinct disease settings rather than a single organ system, which is important when evaluating whether a representation behaves like a reusable foundation model rather than a disease-specific predictor [1, 2].

### 4.2 Architecture

HistoMethyl comprises an image branch and a methylation branch. The image branch uses GigaPath [3] or Feather [17] to encode tissue tiles derived from the whole-slide image. The tile encoder was kept frozen, whereas the slide encoder and image projection head were trainable during pre-training; together, these modules aggregated tile embeddings into a slide-level representation and projected it into the shared embedding space. The methylation branch uses CpGPT [18] to encode the paired DNA methylation profile, represented as CpG-site identifiers with their corresponding beta values, followed by a projection layer that yields a methylation embedding. During pre-training, HistoMethyl concentrates trainable capacity in the slide encoder and projection layers while preserving pre-trained low-level visual features from the frozen tile encoder and using matched DNA methylation profiles as molecular supervision.

Let *u_i_* ∈ ℝ*^d^* and *v_i_* ∈ ℝ*^d^* denote the image and methylation embeddings for case *i*. These embeddings are first compared using the similarity

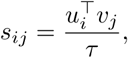

where *τ* is a temperature parameter. We then align them using a SigLIP [38] loss. For a minibatch of *N* matched slide-methylation pairs, the loss is written as

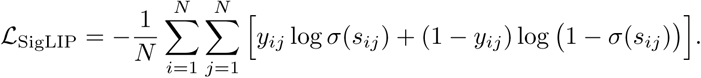

Here, *y_ij_* = 1 for matched slide-methylation pairs and 0 otherwise, and *σ* denotes the sigmoid function. This objective encourages matched slide and methylation embeddings to be close in the shared space while pushing unmatched pairs apart [39].

Although HistoMethyl is trained with paired image and methylation data, it is used as an image-only foundation model at inference. After pre-training, the methylation branch is discarded, and only the projected slide embedding from the image branch is retained for downstream analysis. For mutation prediction, receptor-associated classification, morphology, and survival analysis, we extracted the pre-trained slide embedding after the image-side MLP and used it as input to task-specific supervised models under five-fold cross-validation. Baseline models were evaluated in the same way, using slide embeddings from their respective pre-trained image encoders with matched downstream task models, splits, and preprocessing. All classification tasks were evaluated across three independent runs with random seeds 42, 43, and 44. For DNA methylation beta-value prediction, the regression head and the slide encoder were trained jointly because of the large number of CpG sites and task complexity. In this way, HistoMethyl uses methylation supervision during pre-training to shape the slide representation, while retaining a practical image-only deployment path for downstream clinical and genomic tasks.

### 4.3 Downstream tasks

We selected four downstream task families that test the main qualities expected from a clinically useful pathology foundation model: molecular biomarker inference, diagnostic phenotype recognition, prognostic prediction, and non-image molecular signal recovery from H&E slides [1, 40, 41]. Recent reviews of digital pathology consistently identify molecular prediction, morphology-focused classification, and outcome prediction as central clinical endpoints for whole-slide models [42, 43], while newer multimodal pathology studies increasingly evaluate whether image embeddings can recover molecular information beyond conventional visual tasks [44, 45]. We therefore evaluated HistoMethyl on gene mutation prediction, morphology prediction, survival analysis, and DNA methylation beta-value prediction using the same image-only deployment pathway, so that the downstream benchmark directly tested whether methylation-aware pre-training improved the usefulness of the slide representation.

#### 4.3.1 Gene mutation prediction

Gene mutation prediction was formulated as a set of cohort-specific binary classification tasks. This task is one of the most established molecular benchmarks in computational pathology and directly tests whether the slide embedding captures disease-relevant molecular state rather than only broad appearance [1, 46]. We used pre-specified mutation panels rather than selecting genes after training, and these panels were built around recurrent, clinically interpretable driver genes for each disease context. In glioma-related cohorts, this meant emphasizing canonical diffuse glioma and GBM alterations such as IDH1, ATRX, CIC, FUBP1, EGFR, PTEN, NF1, and RB1, which define major disease subgroups and tumor biology [47, 48]. In colorectal cohorts, the panel was centered on recurrent drivers such as APC, TP53, KRAS, PIK3CA, FBXW7, BRAF, and SMAD4, which remain core genomic events in colorectal tumorigenesis [49, 50]. Consistent with prior work, we attached a small MLP classifier to the pre-trained slide embedding for each mutation and trained it under cohort-specific five-fold cross-validation [3]. The same adaptation protocol was used for the image-only baseline. AUROC was the primary metric reported in the main text, while AUPRC and F1 were retained for the Supplementary Information.

#### 4.3.2 Morphology

Morphology prediction was used as a diagnostic phenotype benchmark to evaluate whether methylation-aligned pre-training preserves the visual information required for conventional histopathologic subtype recognition [28, 51]. Following common whole-slide evaluation practice in computational pathology, we formulated this task as cohort-specific slide-level classification from H&E images, using the two most common morphologic subtypes within each eligible cohort to reduce instability from rare labels and small class sizes [3, 52]. Models were evaluated under the same five-fold cross-validation protocol used for the other classification tasks, with fixed slide embeddings from each pre-trained encoder and the same downstream classifier applied to all baselines and HistoMethyl variants. AUROC was used as the primary metric.

#### 4.3.3 Survival analysis

Prognosis is a core clinical endpoint for digital pathology and evaluates a broader property of the model than a single biomarker task. Whole-slide survival models are increasingly used to assess whether a representation captures global disease severity rather than only diagnostic morphology [28, 53]. This task is also directly relevant to methylation-aware representation learning because DNA methylation biomarkers are widely studied in risk stratification, disease progression, and clinical prognosis across cancer types [54]. For each cohort, we used the HistoMethyl slide embedding as input to a Cox proportional hazards model [55] trained under five-fold cross-validation on right-censored overall survival data. Performance was summarized using C-index, which quantifies the extent to which higher predicted risk scores correspond to earlier observed events among comparable patient pairs [56].

#### 4.3.4 DNA methylation beta-value prediction

DNA methylation beta-value prediction was included as a direct epigenomic recovery benchmark, motivated by prior studies showing that histopathology images can reflect tumor methylation patterns and can be used to infer methylation-based tumor classes or methylation profiles [57–59]. DNA methylation beta values are continuous CpG-level measurements ranging from 0 to 1, reported by methylation array pipelines as the methylated probe intensity normalized by total probe intensity. CpG sites were selected from CpGPT vocabulary sites with sufficient observation rate and beta-value variability, so that site selection did not use held-out labels. Performance was evaluated under cohort-specific five-fold cross-validation using flattened Pearson correlation across all observed CpG-sample pairs, site-wise Pearson correlation to assess per-CpG recovery, and error metrics including MAE to quantify beta-value calibration.

### 4.4 Implementation details

#### Pre-training

HistoMethyl was trained separately with the GigaPath [3] and Feather [17] image backbones. For both variants, the pre-trained tile encoder was frozen and tissue tiles were aggregated by the corresponding slide encoder. The slide encoder and image projection head were optimized during methylation alignment, while the CpGPT methylation encoder was kept frozen and followed by a trainable methylation projection head. Image and methylation embeddings were projected into a shared space (768 for the GigaPath-based variant and 256 for the Feather-based variant) with MLP projection heads, and paired slide–methylation representations were aligned with the SigLIP objective using an initial temperature of 0.07.

Models were optimized with AdamW using a learning rate of 1×10^−4^, weight decay of 0.01, betas of 0.9 and 0.95, gradient clipping of 5, and mixed bfloat16 precision. The batch size was 48 for GigaPath and 128 for Feather. Training ran for up to 30 epochs for the GigaPath backbone and up to 100 epochs for the Feather backbone, with validation loss used for checkpointing and early stopping. Pre-training was performed on one node with four NVIDIA H100 GPUs using distributed data parallel training.

#### Downstream task adaptation

For downstream classification tasks, pre-trained slide embeddings were extracted from the image branch and evaluated with the same supervised adaptation protocol for HistoMethyl and the matched image-only baselines. Mutation, receptor, and morphology tasks used a one-hidden-layer MLP classifier with hidden dimension 256, batch size 64, learning rate 3 × 10^−4^, and 50 training epochs. Classification tasks were evaluated with cohort-specific five-fold cross-validation. Survival prediction used the same fixed slide embeddings as input to a Cox proportional hazards model and was evaluated by concordance index.

For DNA methylation beta-value prediction, CpG targets were selected within each training fold from CpGPT vocabulary sites with sufficient observation rate and beta-value variability. A multi-output regression head was trained to predict beta values from the slide representation using masked mean squared error so that missing CpG measurements did not contribute to the loss. The slide encoder was fine-tuned, and the regression head was optimized with AdamW, batch size 64, learning rate 2 × 10^−4^, and backbone-specific training schedules of 30 epochs for models using the GigaPath backbone and 70 epochs for models using the Feather backbone. Prediction performance was summarized using global Pearson correlation, site-wise Pearson correlation, and error metrics on held-out folds.

## Supporting information

Supplementary Infomation

## 5 Data availability

The TCGA [14] dataset is available at https://www.cancer.gov/ccg/research/genome-sequencing/tcga. The CPTAC dataset is available at https://portal.gdc.cancer.gov/projects/CPTAC-3. The SurGen dataset is available at https://github.com/CraigMyles/SurGen-Dataset.

## 6 Code availability

Our project code is available at https://github.com/wangd12rpi/histomethyl.

## References

[1] Chen, R.J., Ding, T., Lu, M.Y., Williamson, D.F., Jaume, G., Song, A.H., Chen, B., Zhang, A., Shao, D., Shaban, M., et al.: Towards a general-purpose foundation model for computational pathology. Nature medicine 30(3), 850–862 (2024)

[2] Ding, T., Wagner, S.J., Song, A.H., Chen, R.J., Lu, M.Y., Zhang, A., Vaidya, A.J., Jaume, G., Shaban, M., Kim, A., et al.: A multimodal whole-slide foundation model for pathology. Nature Medicine, 1–13 (2025)

[3] Xu, H., Usuyama, N., Bagga, J., Zhang, S., Rao, R., Naumann, T., Wong, C., Gero, Z., González, J., Gu, Y., Xu, Y., Wei, M., Wang, W., Ma, S., Wei, F., Yang, J., Li, C., Gao, J., Rosemon, J., Bower, T., Lee, S., Weerasinghe, R., Wright, B.J., Robicsek, A., Piening, B., Bifulco, C., Wang, S., Poon, H.: A whole-slide foundation model for digital pathology from real-world data. Nature (2024)

[4] Chen, W., Zhang, P., Tran, T.N., Xiao, Y., Li, S., Shah, V.V., Cheng, H., Brannan, K.W., Youker, K., Lai, L., et al.: A visual–omics foundation model to bridge histopathology with spatial transcriptomics. Nature Methods 22(7), 1568–1582 (2025)

[5] Huang, T., Liu, T., Babadi, M., Ying, R., Jin, W.: Stpath: a generative foundation model for integrating spatial transcriptomics and whole-slide images. NPJ Digital Medicine 8(1), 659 (2025)

[6] Jaume, G., Oldenburg, L., Vaidya, A., Chen, R.J., Williamson, D.F.K., Peeters, T., Song, A.H., Mahmood, F.: Transcriptomics-guided slide representation learning in computational pathology. In: Proceedings of the IEEE/CVF Conference on Computer Vision and Pattern Recognition (CVPR), pp. 9632–9644 (2024)

[7] Gindra, R.H., Palla, G., Nguyen, M., Wagner, S.J., Tran, M., Theis, F.J., Saur, D., Crawford, L., Peng, T.: A Large-Scale Benchmark of Cross-Modal Learning for Histology and Gene Expression in Spatial Transcriptomics (2025). https://arxiv.org/abs/2508.01490

[8] Kather, J.N., Pearson, A.T., Halama, N., Jäger, D., Krause, J., Loosen, S.H., Marx, A., Boor, P., Tacke, F., Neumann, U.P., et al.: Deep learning can predict microsatellite instability directly from histology in gastrointestinal cancer. Nature medicine 25(7), 1054–1056 (2019)

[9] Liang, W.-W., Lu, R.J.-H., Jayasinghe, R.G., Foltz, S.M., Porta-Pardo, E., Geffen, Y., Wendl, M.C., Lazcano, R., Kolodziejczak, I., Song, Y., et al.: Integrative multi-omic cancer profiling reveals dna methylation patterns associated with therapeutic vulnerability and cell-of-origin. Cancer cell 41(9), 1567–1585 (2023)

[10] Capper, D., Jones, D.T., Sill, M., Hovestadt, V., Schrimpf, D., Sturm, D., Koelsche, C., Sahm, F., Chavez, L., Reuss, D.E., et al.: Dna methylation-based classification of central nervous system tumours. Nature 555(7697), 469–474 (2018)

[11] Louis, D.N., Perry, A., Wesseling, P., Brat, D.J., Cree, I.A., Figarella-Branger, D., Hawkins, C., Ng, H., Pfister, S.M., Reifenberger, G., et al.: The 2021 who classification of tumors of the central nervous system: a summary. Neuro-oncology 23(8), 1231–1251 (2021)

[12] Jaunmuktane, Z., Capper, D., Jones, D.T., Schrimpf, D., Sill, M., Dutt, M., Suraweera, N., Pfister, S.M., Deimling, A., Brandner, S.: Methylation array profiling of adult brain tumours: diagnostic outcomes in a large, single centre. Acta neuropathologica communications 7(1), 24 (2019)

[13] Patel, A., Göbel, K., Ille, S., Hinz, F., Schoebe, N., Bogumil, H., Meyer, J., Brehm, M., Kardo, H., Schrimpf, D., et al.: Prospective, multicenter validation of a platform for rapid molecular profiling of central nervous system tumors. Nature medicine 31(5), 1567–1577 (2025)

[14] Tomczak, K., Czerwińska, P., Wiznerowicz, M.: Review the cancer genome atlas (tcga): an immeasurable source of knowledge. Contemporary Oncology/Współczesna Onkologia 2015(1), 68–77 (2015)

[15] Consortium, N.C.I.C.P.T.A., et al.: The clinical proteomic tumor analysis consortium glioblastoma multiforme collection (cptac-gbm). The Cancer Imaging Archive (2023)

[16] Myles, C., Um, I.H., Marshall, C., Harris-Birtill, D., Harrison, D.J.: Surgen: 1020 h&e-stained whole-slide images with survival and genetic markers. GigaScience 14, 086 (2025)

[17] Shao, D., Chen, R.J., Song, A.H., Runevic, J., Lu, M.Y., Ding, T.,, Mahmood, F.: Do multiple instance learning models transfer? In: International Conference on Machine Learning (2025)

[18] Lima Camillo, L.P., Sehgal, R., Armstrong, J., Miller, H.E., Lasky-Su, J.A., Higgins-Chen, A.T., Horvath, S., Wang, B.: Cpgpt: a foundation model for dna methylation. bioRxiv, 2024–10 (2024)

[19] Saghafinia, S., Mina, M., Riggi, N., Hanahan, D., Ciriello, G.: Pan-cancer landscape of aberrant dna methylation across human tumors. Cell reports 25(4), 1066–1080 (2018)

[20] Fu, Y., Jung, A.W., Torne, R.V., Gonzalez, S., Vöhringer, H., Shmatko, A., Yates, L.R., Jimenez-Linan, M., Moore, L., Gerstung, M.: Pan-cancer computational histopathology reveals mutations, tumor composition and prognosis. Nature cancer 1(8), 800–810 (2020)

[21] Wulczyn, E., Steiner, D.F., Xu, Z., Sadhwani, A., Wang, H., Flament-Auvigne, I., Mermel, C.H., Chen, P.-H.C., Liu, Y., Stumpe, M.C.: Deep learning-based survival prediction for multiple cancer types using histopathology images. PloS one 15(6), 0233678 (2020)

[22] Zeiler, M.D., Fergus, R.: Visualizing and understanding convolutional networks. In: European Conference on Computer Vision, pp. 818–833 (2014). Springer

[23] Kers, J., Bülow, R.D., Klinkhammer, B.M., Breimer, G.E., Fontana, F., Abiola, A.A., Hofstraat, R., Corthals, G.L., Peters-Sengers, H., Djudjaj, S., et al.: Deep learning-based classification of kidney transplant pathology: a retrospective, multicentre, proof-of-concept study. The Lancet Digital Health 4(1), 18–26 (2022)

[24] Eckardt, J.-N., Middeke, J.M., Riechert, S., Schmittmann, T., Sulaiman, A.S., Kramer, M., Sockel, K., Kroschinsky, F., Schuler, U., Schetelig, J., et al.: Deep learning detects acute myeloid leukemia and predicts npm1 mutation status from bone marrow smears. Leukemia 36(1), 111 (2021)

[25] Eckardt, J.-N., Srivastava, I., Schulze, F., Winter, S., Schmittmann, T., Riechert, S., Schneider, M.M., Reichel, L., Gediga, M.E.H., Sockel, K., et al.: Image-based explainable artificial intelligence accurately identifies myelodysplastic neoplasms beyond conventional signs of dysplasia. NPJ Precision Oncology (2025)

[26] Wang, X., Tan, T., Gao, Y., Zhou, H.-Y., Zhang, T., Han, L., Portaluri, A., Marcus, E., Lu, C., Drukker, C., Teuwen, J., Beets-Tan, R., Wang, S., Karssemeijer, N., Mann, R.: Mammo-age: deep learning estimation of breast age from mammograms. Nature Communications 16 (2025) 10.1038/s41467-025-65923-5

[27] Abrol, A., Fu, Z., Salman, M., Silva, R., Du, Y., Plis, S., Calhoun, V.: Deep learning encodes robust discriminative neuroimaging representations to outperform standard machine learning. Nature communications 12(1), 353 (2021)

[28] Lu, M.Y., Williamson, D.F., Chen, T.Y., Chen, R.J., Barbieri, M., Mahmood, F.: Data-efficient and weakly supervised computational pathology on whole-slide images. Nature biomedical engineering 5(6), 555–570 (2021)

[29] Compton, C.C., Robb, J.A., Anderson, M.W., Berry, A.B., Birdsong, G.G., Bloom, K.J., Branton, P.A., Crothers, J.W., Cushman-Vokoun, A.M., Hicks, D.G., et al.: Preanalytics and precision pathology: pathology practices to ensure molecular integrity of cancer patient biospecimens for precision medicine. Archives of pathology & laboratory medicine 143(11), 1346–1363 (2019)

[30] Dagogo-Jack, I., Robinson, H., Mino-Kenudson, M., Farago, A.F., Kamesan, V., Iafrate, A.J., Shaw, A.T., Lennerz, J.K.: Expediting comprehensive molecular analysis to optimize initial treatment of lung cancer patients with minimal smoking history. Journal of Thoracic Oncology 14(5), 835–843 (2019)

[31] Ossowski, S., Neeman, E., Borden, C., Stram, D., Giraldo, L., Kotak, D., Thomas, S., Suga, J.M., Lin, A., Liu, R.: Improving time to molecular testing results in patients with newly diagnosed, metastatic non–small-cell lung cancer. JCO Oncology Practice 18(11), 1874–1884 (2022)

[32] Tamborero, D., Dienstmann, R., Rachid, M.H., Boekel, J., Baird, R., Braña, I., De Petris, L., Yachnin, J., Massard, C., Opdam, F.L., et al.: Support systems to guide clinical decision-making in precision oncology: The cancer core europe molecular tumor board portal. Nature medicine 26(7), 992–994 (2020)

[33] Cooper, L.A., Demicco, E.G., Saltz, J.H., Powell, R.T., Rao, A., Lazar, A.J.: Pancancer insights from the cancer genome atlas: the pathologist’s perspective. The Journal of pathology 244(5), 512–524 (2018)

[34] Gutman, D.A., Cobb, J., Somanna, D., Park, Y., Wang, F., Kurc, T., Saltz, J.H., Brat, D.J., Cooper, L.A., Kong, J.: Cancer digital slide archive: an informatics resource to support integrated in silico analysis of tcga pathology data. Journal of the American Medical Informatics Association 20(6), 1091–1098 (2013)

[35] Novis, D.A., Zarbo, R.J.: Interinstitutional comparison of frozen section turnaround time. Archives of pathology & laboratory medicine 121(6), 559 (1997)

[36] Zakharova, G., Efimov, V., Raevskiy, M., Rumiantsev, P., Gudkov, A., Belogurova-Ovchinnikova, O., Sorokin, M., Buzdin, A.: Reclassification of tcga diffuse glioma profiles linked to transcriptomic, epigenetic, genomic and clinical data, according to the 2021 who cns tumor classification. International journal of molecular sciences 24(1), 157 (2022)

[37] Singh, J., Sahu, S., Mohan, T., Mahajan, S., Sharma, M.C., Sarkar, C., Suri, V.: Current status of dna methylation profiling in neuro-oncology as a diagnostic support tool: A review. Neuro-Oncology Practice 10(6), 518–526 (2023)

[38] Zhai, X., Mustafa, B., Kolesnikov, A., Beyer, L.: Sigmoid loss for language image pre-training. In: Proceedings of the IEEE/CVF International Conference on Computer Vision, pp. 11975–11986 (2023)

[39] Radford, A., Kim, J.W., Hallacy, C., Ramesh, A., Goh, G., Agarwal, S., Sastry, G., Askell, A., Mishkin, P., Clark, J., et al.: Learning transferable visual models from natural language supervision. In: International Conference on Machine Learning, pp. 8748–8763 (2021). PmLR

[40] Wang, X., Zhao, J., Marostica, E., Yuan, W., Jin, J., Zhang, J., Li, R., Tang, H., Wang, K., Li, Y., et al.: A pathology foundation model for cancer diagnosis and prognosis prediction. Nature 634(8035), 970–978 (2024)

[41] Vorontsov, E., Bozkurt, A., Casson, A., Shaikovski, G., Zelechowski, M., Severson, K., Zimmermann, E., Hall, J., Tenenholtz, N., Fusi, N., et al.: A foundation model for clinical-grade computational pathology and rare cancers detection. Nature medicine 30(10), 2924–2935 (2024)

[42] Rad, M.S., Huang, J.V., Hosseini, M.M., Choudhary, R., Siezen, H., Akabari, R., Jamaspishvili, T., El-Zammar, O., Patel, P., Carello, S.J., et al.: Deep learning for digital pathology: A critical overview of methodological framework. Journal of Pathology Informatics, 100514 (2025)

[43] Dang, C., Qi, Z., Xu, T., Gu, M., Chen, J., Wu, J., Lin, Y., Qi, X.: Deep learning–powered whole slide image analysis in cancer pathology. Laboratory Investigation 105(7), 104186 (2025)

[44] Jang, H.-J., Lee, S.H.: Ai-driven digital pathology: Deep learning and multimodal integration for precision oncology. International Journal of Molecular Sciences 27(1), 379 (2025)

[45] Nateghi, R., Sun, A., Dang, H., Handa, N., Schnauss, M., Michael, J., Jang, J.W., Nezami, B.G., Saft, M., Neill, C., et al.: Prediction of molecular subtypes from histology: Ai-driven analysis of prostate cancer morphological patterns and therapeutic implications. npj Precision Oncology (2026)

[46] Coudray, N., Ocampo, P.S., Sakellaropoulos, T., Narula, N., Snuderl, M., Fenyö, D., Moreira, A.L., Razavian, N., Tsirigos, A.: Classification and mutation prediction from non–small cell lung cancer histopathology images using deep learning. Nature medicine 24(10), 1559–1567 (2018)

[47] Jiao, Y., Killela, P.J., Reitman, Z.J., Rasheed, B.A., Heaphy, C.M., De Wilde, R.F., Rodriguez, F.J., Rosemberg, S., Oba-Shinjo, S.M., Marie, S.K.N., et al.: Frequent atrx, cic, fubp1 and idh1 mutations refine the classification of malignant gliomas. Oncotarget 3(7), 709 (2012)

[48] Kinnersley, B., Jung, J., Cornish, A.J., Chubb, D., Laxton, R., Frangou, A., Gruber, A.J., Sud, A., Caravagna, G., Sottoriva, A., et al.: Genomic landscape of diffuse glioma revealed by whole genome sequencing. Nature communications 16(1), 4233 (2025)

[49] Navarro-Jiménez, M., González, B., Mulet, N., Hierro, C., Alonso, S.: Kras-targeted therapies in colorectal cancer: a systematic analysis of mutations, inhibitors, and clinical trials. NPJ Precision Oncology 9(1), 380 (2025)

[50] Su, Z., El Hage, M., Linnebahcher, M.: Mutation patterns in colorectal cancer and their relationship with prognosis. Heliyon 10(17) (2024)

[51] Chelebian, E., Avenel, C., Wählby, C.: Combining spatial transcriptomics with tissue morphology. Nature Communications 16(1), 4452 (2025)

[52] Noorbakhsh, J., Farahmand, S., Foroughi Pour, A., Namburi, S., Caruana, D., Rimm, D., Soltanieh-Ha, M., Zarringhalam, K., Chuang, J.H.: Deep learning-based cross-classifications reveal conserved spatial behaviors within tumor histological images. Nature communications 11(1), 6367 (2020)

53. [53] Mobadersany, P., Yousefi, S., Amgad, M., Gutman, D.A., Barnholtz-Sloan, J.S., Velázquez Vega, J.E., Brat, D.J., Cooper, L.A.: Predicting cancer outcomes from histology and genomics using convolutional networks. Proceedings of the National Academy of Sciences 115(13), 2970–2979 (2018)

[54] Fan, Z., Edelmann, D., Yuan, T., Köhler, B.C., Hoffmeister, M., Brenner, H.: Developing survival prediction models in colorectal cancer using epigenome-wide dna methylation data from whole blood. NPJ Precision Oncology 8(1), 191 (2024)

[55] Cox, D.R. Regression models and life-tables. Journal of the royal statistical society: Series B (methodological) 34(2), 187–202 (1972)

[56] Liu, H., Kurc, T.: Deep learning for survival analysis in breast cancer with whole slide image data. Bioinformatics 38(14), 3629–3637 (2022)

[57] Zheng, H., Momeni, A., Cedoz, P.-L., Vogel, H., Gevaert, O.: Whole slide images reflect dna methylation patterns of human tumors. NPJ genomic medicine 5(1), 11 (2020)

[58] Hoang, D.-T., Shulman, E.D., Turakulov, R., Abdullaev, Z., Singh, O., Campagnolo, E.M., Lalchungnunga, H., Stone, E.A., Nasrallah, M.P., Ruppin, E., et al.: Prediction of dna methylation-based tumor types from histopathology in central nervous system tumors with deep learning. Nature Medicine 30(7), 1952–1961 (2024)

[59] Hoang, D.-T., Shulman, E.D., Dhruba, S.R., Nair, N.U., Barman, R.K., Cantore, T., Biswas, S., Lalchungnunga, H., Singh, O., Chung, Y., et al.: Path2omics enhances transcriptomic and methylation prediction accuracy from tumor histopathology. Cancer research 85(24), 5098–5112 (2025)

