## Supplementary Infomation for "A multimodal foundation model linking histopathology and DNA methylation"

### Supplementary information

### 1 Detailed results

#### 1.1 Complete downstream benchmark tables

**Supplementary Table 1**: Mutation prediction performance across filtered cancer–gene tasks. Cells report AUROC / AUPRC, with cross-validation standard deviations shown below each metric. Cancer-specific blocks correspond to the filtered mutation tasks used for the main barplot calculations. Bold values indicate the best model for each metric within a task, and underlined values indicate the second-best model.

| Task | GigaPath | HistoMethyl GigaPath | Feather | HistoMethyl Feather |
|---|---|---|---|---|
| **BRCA** (AUROC / AUPRC) | | | | |
| AKT1 | 0.586 / 0.072<br>±0.211/±0.057 | **0.699** / **0.091**<br>±0.162/±0.049 | 0.566 / 0.056<br>±0.174/±0.025 | 0.584 / 0.063<br>±0.180/±0.048 |
| CDH1 | 0.840 / 0.410<br>±0.054/±0.102 | 0.841 / 0.379<br>±0.075/±0.111 | 0.833 / **0.431**<br>±0.068/±0.134 | **0.848** / 0.378<br>±0.047/±0.110 |
| GATA3 | 0.534 / 0.194<br>±0.087/±0.080 | **0.599** / **0.201**<br>±0.071/±0.100 | 0.570 / 0.191<br>±0.081/±0.071 | 0.544 / 0.183<br>±0.127/±0.090 |
| KMT2C | 0.539 / 0.116<br>±0.118/±0.037 | **0.575** / **0.136**<br>±0.083/±0.029 | 0.551 / 0.107<br>±0.070/±0.023 | 0.556 / 0.105<br>±0.060/±0.019 |
| MAP3K1 | 0.678 / 0.227<br>±0.082/±0.091 | 0.725 / 0.240<br>±0.120/±0.129 | 0.596 / 0.145<br>±0.123/±0.054 | **0.782** / **0.319**<br>±0.099/±0.102 |
| PIK3CA | 0.670 / 0.461<br>±0.046/±0.051 | 0.671 / 0.505<br>±0.049/±0.066 | 0.670 / 0.485<br>±0.050/±0.079 | **0.692** / **0.519**<br>±0.054/±0.063 |
| PTEN | 0.466 / 0.057<br>±0.129/±0.017 | **0.580** / 0.106<br>±0.124/±0.088 | 0.454 / 0.070<br>±0.164/±0.035 | 0.534 / **0.160**<br>±0.146/±0.128 |
| TP53 | 0.785 / 0.677<br>±0.054/±0.070 | 0.776 / 0.669<br>±0.058/±0.083 | 0.799 / 0.678<br>±0.040/±0.065 | **0.818** / **0.701**<br>±0.041/±0.054 |
| **COAD** (AUROC / AUPRC) | | | | |
| APC | 0.614 / 0.724<br>±0.070/±0.064 | 0.638 / 0.744<br>±0.087/±0.072 | 0.640 / 0.727<br>±0.087/±0.071 | **0.724** / **0.807**<br>±0.069/±0.064 |
| BRAF | 0.674 / 0.364<br>±0.151/±0.162 | 0.671 / 0.385<br>±0.136/±0.211 | **0.706** / **0.414**<br>±0.143/±0.163 | 0.697 / 0.405<br>±0.136/±0.169 |
| FBXW7 | 0.490 / **0.276**<br>±0.106/±0.144 | **0.553** / 0.276<br>±0.143/±0.126 | 0.476 / 0.211<br>±0.139/±0.065 | 0.530 / 0.254<br>±0.101/±0.065 |
| KRAS | 0.538 / 0.435<br>±0.058/±0.055 | 0.466 / 0.404<br>±0.092/±0.076 | 0.514 / 0.449<br>±0.105/±0.096 | **0.583** / **0.477**<br>±0.101/±0.092 |
| PIK3CA | 0.569 / 0.340<br>±0.102/±0.086 | **0.596** / **0.376**<br>±0.099/±0.128 | 0.586 / 0.345<br>±0.091/±0.075 | 0.581 / 0.373<br>±0.072/±0.070 |
| SMAD4 | **0.639** / **0.321**<br>±0.115/±0.149 | 0.608 / 0.234<br>±0.119/±0.096 | 0.507 / 0.163<br>±0.144/±0.064 | 0.598 / 0.219<br>±0.136/±0.079 |

Continued on next page

**Supplementary Table 1**: Mutation prediction performance across filtered cancer–gene tasks (continued)

| Task | GigaPath | HistoMethyl GigaPath | Feather | HistoMethyl Feather |
|---|---|---|---|---|
| TP53 | 0.613 / 0.620 ±0.087/±0.085 | 0.569 / 0.585 ±0.094/±0.088 | 0.630 / 0.638 ±0.090/±0.094 | 0.600 / 0.605 ±0.113/±0.102 |
| **GBM** (AUROC / AUPRC) | | | | |
| EGFR | 0.489 / 0.194 ±0.088/±0.044 | 0.488 / 0.223 ±0.098/±0.083 | 0.546 / 0.244 ±0.062/±0.046 | 0.529 / 0.231 ±0.100/±0.051 |
| NF1 | 0.621 / 0.200 ±0.125/±0.117 | 0.634 / 0.170 ±0.124/±0.051 | 0.577 / 0.138 ±0.107/±0.051 | 0.670 / 0.222 ±0.118/±0.104 |
| PIK3CA | 0.417 / 0.174 ±0.130/±0.100 | 0.609 / 0.244 ±0.148/±0.171 | 0.595 / 0.180 ±0.118/±0.088 | 0.526 / 0.151 ±0.171/±0.085 |
| PIK3R1 | 0.552 / 0.134 ±0.121/±0.046 | 0.583 / 0.141 ±0.109/±0.043 | 0.453 / 0.101 ±0.106/±0.039 | 0.597 / 0.149 ±0.136/±0.052 |
| PTEN | 0.602 / 0.324 ±0.074/±0.080 | 0.635 / 0.330 ±0.059/±0.059 | 0.547 / 0.279 ±0.067/±0.057 | 0.639 / 0.355 ±0.066/±0.083 |
| RB1 | 0.452 / 0.110 ±0.144/±0.069 | 0.596 / 0.173 ±0.136/±0.106 | 0.425 / 0.132 ±0.123/±0.111 | 0.597 / 0.204 ±0.135/±0.127 |
| TP53 | 0.673 / 0.455 ±0.053/±0.096 | 0.654 / 0.444 ±0.079/±0.101 | 0.623 / 0.418 ±0.097/±0.113 | 0.676 / 0.474 ±0.072/±0.086 |
| **HNSC** (AUROC / AUPRC) | | | | |
| CASP8 | 0.642 / 0.245 ±0.068/±0.089 | 0.652 / 0.234 ±0.081/±0.067 | 0.431 / 0.122 ±0.093/±0.023 | 0.722 / 0.291 ±0.091/±0.129 |
| CDKN2A | 0.509 / 0.226 ±0.097/±0.066 | 0.618 / 0.290 ±0.120/±0.100 | 0.513 / 0.199 ±0.130/±0.054 | 0.585 / 0.247 ±0.089/±0.056 |
| FAT1 | 0.437 / 0.199 ±0.110/±0.043 | 0.457 / 0.214 ±0.081/±0.044 | 0.466 / 0.232 ±0.096/±0.066 | 0.471 / 0.203 ±0.068/±0.027 |
| HRAS | 0.382 / 0.074 ±0.094/±0.014 | 0.556 / 0.125 ±0.163/±0.065 | 0.455 / 0.087 ±0.100/±0.017 | 0.560 / 0.121 ±0.111/±0.034 |
| NOTCH1 | 0.445 / 0.221 ±0.101/±0.062 | 0.538 / 0.237 ±0.083/±0.047 | 0.385 / 0.167 ±0.099/±0.034 | 0.613 / 0.280 ±0.102/±0.070 |
| PIK3CA | 0.493 / 0.247 ±0.093/±0.062 | 0.502 / 0.221 ±0.088/±0.054 | 0.388 / 0.177 ±0.095/±0.053 | 0.487 / 0.225 ±0.118/±0.105 |
| TP53 | 0.702 / 0.824 ±0.083/±0.058 | 0.657 / 0.792 ±0.096/±0.070 | 0.723 / 0.818 ±0.074/±0.060 | 0.698 / 0.813 ±0.079/±0.054 |
| **KIRC** (AUROC / AUPRC) | | | | |
| BAP1 | 0.649 / 0.167 ±0.137/±0.077 | 0.667 / 0.197 ±0.117/±0.103 | 0.526 / 0.130 ±0.111/±0.074 | 0.685 / 0.218 ±0.136/±0.118 |
| KDM5C | 0.504 / 0.122 ±0.186/±0.100 | 0.602 / 0.151 ±0.140/±0.124 | 0.714 / 0.163 ±0.132/±0.116 | 0.465 / 0.065 ±0.190/±0.054 |
| MTOR | 0.489 / 0.105 ±0.102/±0.068 | 0.600 / 0.189 ±0.092/±0.130 | 0.463 / 0.057 ±0.119/±0.018 | 0.480 / 0.106 ±0.157/±0.093 |
| PBRM1 | 0.548 / 0.345 ±0.068/±0.060 | 0.564 / 0.358 ±0.057/±0.042 | 0.587 / 0.370 ±0.060/±0.047 | 0.557 / 0.360 ±0.050/±0.044 |
| SETD2 | 0.563 / 0.150 ±0.102/±0.069 | 0.611 / 0.182 ±0.100/±0.060 | 0.619 / 0.150 ±0.113/±0.050 | 0.663 / 0.200 ±0.069/±0.112 |

Continued on next page

**Supplementary Table 1**: Mutation prediction performance across filtered cancer–gene tasks (continued)

| Task | GigaPath | HistoMethyl GigaPath | Feather | HistoMethyl Feather |
|---|---|---|---|---|
| VHL | 0.550 / 0.407 ±0.041/±0.046 | 0.554 / 0.417 ±0.051/±0.056 | 0.572 / 0.412 ±0.046/±0.041 | 0.550 / 0.424 ±0.044/±0.051 |
| **KIRP** (AUROC / AUPRC) | | | | |
| KMT2C | 0.522 / 0.123 ±0.167/±0.059 | 0.612 / 0.204 ±0.205/±0.232 | 0.548 / 0.142 ±0.237/±0.096 | 0.625 / 0.235 ±0.230/±0.207 |
| KMT2D | 0.533 / 0.162 ±0.192/±0.097 | 0.551 / 0.185 ±0.147/±0.177 | 0.464 / 0.155 ±0.222/±0.086 | 0.558 / 0.210 ±0.185/±0.172 |
| MET | 0.551 / 0.182 ±0.171/±0.172 | 0.539 / 0.126 ±0.146/±0.053 | 0.408 / 0.160 ±0.244/±0.169 | 0.507 / 0.147 ±0.215/±0.109 |
| SETD2 | 0.636 / 0.242 ±0.172/±0.186 | 0.695 / 0.186 ±0.126/±0.144 | 0.442 / 0.111 ±0.239/±0.083 | 0.577 / 0.129 ±0.202/±0.064 |
| **LGG** (AUROC / AUPRC) | | | | |
| ATRX | 0.755 / 0.710 ±0.098/±0.119 | 0.718 / 0.633 ±0.073/±0.083 | 0.693 / 0.622 ±0.071/±0.101 | 0.659 / 0.592 ±0.063/±0.088 |
| EGFR | 0.570 / 0.154 ±0.222/±0.111 | 0.622 / 0.166 ±0.248/±0.147 | 0.439 / 0.125 ±0.115/±0.133 | 0.546 / 0.198 ±0.256/±0.208 |
| FUBP1 | 0.672 / 0.196 ±0.105/±0.071 | 0.629 / 0.251 ±0.128/±0.175 | 0.518 / 0.224 ±0.139/±0.135 | 0.554 / 0.198 ±0.201/±0.118 |
| IDH1 | 0.677 / 0.862 ±0.090/±0.046 | 0.712 / 0.879 ±0.068/±0.041 | 0.570 / 0.801 ±0.071/±0.050 | 0.644 / 0.843 ±0.080/±0.051 |
| NF1 | 0.560 / 0.194 ±0.176/±0.189 | 0.650 / 0.284 ±0.160/±0.206 | 0.365 / 0.081 ±0.176/±0.048 | 0.565 / 0.128 ±0.152/±0.062 |
| TP53 | 0.790 / 0.816 ±0.053/±0.040 | 0.776 / 0.791 ±0.033/±0.035 | 0.689 / 0.685 ±0.064/±0.073 | 0.677 / 0.692 ±0.051/±0.055 |
| **LUAD** (AUROC / AUPRC) | | | | |
| BRAF | 0.475 / 0.093 ±0.084/±0.021 | 0.562 / 0.134 ±0.132/±0.052 | 0.466 / 0.177 ±0.173/±0.137 | 0.563 / 0.153 ±0.139/±0.092 |
| EGFR | 0.662 / 0.284 ±0.112/±0.126 | 0.550 / 0.189 ±0.081/±0.061 | 0.496 / 0.185 ±0.130/±0.097 | 0.602 / 0.229 ±0.078/±0.091 |
| NF1 | 0.636 / 0.264 ±0.082/±0.092 | 0.508 / 0.163 ±0.119/±0.082 | 0.463 / 0.172 ±0.055/±0.108 | 0.616 / 0.208 ±0.093/±0.067 |
| SMARCA4 | 0.660 / 0.329 ±0.096/±0.094 | 0.693 / 0.267 ±0.082/±0.124 | 0.554 / 0.225 ±0.112/±0.091 | 0.641 / 0.297 ±0.105/±0.120 |
| TP53 | 0.740 / 0.683 ±0.055/±0.068 | 0.715 / 0.650 ±0.043/±0.067 | 0.723 / 0.657 ±0.040/±0.053 | 0.716 / 0.643 ±0.036/±0.049 |
| **LUSC** (AUROC / AUPRC) | | | | |
| CDKN2A | 0.609 / 0.448 ±0.134/±0.146 | 0.631 / 0.365 ±0.091/±0.093 | 0.563 / 0.254 ±0.130/±0.082 | 0.541 / 0.236 ±0.160/±0.080 |
| FAT1 | 0.536 / 0.219 ±0.152/±0.073 | 0.504 / 0.179 ±0.072/±0.041 | 0.467 / 0.221 ±0.134/±0.115 | 0.477 / 0.189 ±0.141/±0.083 |
| KMT2D | 0.463 / 0.330 ±0.127/±0.108 | 0.415 / 0.295 ±0.124/±0.124 | 0.489 / 0.316 ±0.127/±0.105 | 0.477 / 0.303 ±0.103/±0.087 |

Continued on next page

**Supplementary Table 1**: Mutation prediction performance across filtered cancer–gene tasks (continued)

| Task | GigaPath | HistoMethyl GigaPath | Feather | HistoMethyl Feather |
|---|---|---|---|---|
| NOTCH1 | 0.561 / 0.192 ±0.190/±0.144 | **0.635** / 0.180 ±0.148/±0.077 | 0.381 / 0.096 ±0.137/±0.028 | 0.569 / **0.249** ±0.175/±0.152 |
| PIK3CA | 0.353 / 0.117 ±0.116/±0.034 | 0.416 / 0.129 ±0.116/±0.033 | **0.535** / **0.236** ±0.084/±0.125 | 0.347 / 0.122 ±0.125/±0.043 |
| PTEN | 0.710 / 0.242 ±0.126/±0.079 | **0.724** / **0.296** ±0.117/±0.135 | 0.531 / 0.227 ±0.203/±0.152 | 0.493 / 0.145 ±0.160/±0.056 |
| RB1 | 0.597 / 0.143 ±0.218/±0.081 | 0.470 / 0.182 ±0.294/±0.206 | 0.611 / 0.193 ±0.274/±0.128 | **0.686** / **0.250** ±0.174/±0.179 |
| TP53 | 0.412 / 0.835 ±0.094/±0.043 | 0.410 / 0.835 ±0.128/±0.051 | 0.521 / 0.868 ±0.195/±0.065 | **0.599** / **0.901** ±0.137/±0.043 |
| **READ** (AUROC / AUPRC) | | | | |
| APC | 0.796 / 0.909 ±0.095/±0.052 | 0.821 / **0.912** ±0.112/±0.062 | 0.688 / 0.806 ±0.157/±0.114 | **0.832** / 0.911 ±0.108/±0.064 |
| FBXW7 | 0.375 / 0.266 ±0.234/±0.192 | 0.263 / 0.204 ±0.223/±0.172 | **0.447** / 0.191 ±0.177/±0.052 | 0.382 / **0.275** ±0.231/±0.226 |
| KRAS | 0.469 / 0.391 ±0.130/±0.105 | 0.483 / 0.414 ±0.140/±0.117 | **0.487** / **0.436** ±0.154/±0.124 | 0.462 / 0.424 ±0.107/±0.099 |
| NRAS | 0.304 / 0.112 ±0.197/±0.036 | **0.358** / **0.119** ±0.180/±0.031 | 0.303 / 0.117 ±0.260/±0.045 | 0.246 / 0.106 ±0.212/±0.041 |
| SMAD4 | **0.588** / **0.375** ±0.217/±0.188 | 0.477 / 0.340 ±0.210/±0.211 | 0.533 / 0.289 ±0.191/±0.129 | 0.566 / 0.312 ±0.172/±0.151 |
| TP53 | 0.740 / 0.785 ±0.138/±0.112 | 0.743 / 0.778 ±0.147/±0.126 | 0.722 / 0.753 ±0.122/±0.120 | **0.758** / **0.798** ±0.133/±0.112 |
| **SR386-READ** (AUROC / AUPRC) | | | | |
| RAS | 0.517 / 0.439 ±0.081/±0.079 | 0.494 / 0.448 ±0.114/±0.122 | 0.502 / 0.421 ±0.125/±0.095 | **0.559** / **0.488** ±0.106/±0.089 |
| **STAD** (AUROC / AUPRC) | | | | |
| ARID1A | 0.513 / 0.353 ±0.112/±0.092 | **0.549** / 0.340 ±0.089/±0.052 | 0.525 / **0.366** ±0.080/±0.062 | 0.481 / 0.309 ±0.086/±0.048 |
| CDH1 | 0.549 / 0.148 ±0.179/±0.136 | 0.422 / 0.089 ±0.199/±0.037 | 0.310 / 0.069 ±0.177/±0.018 | **0.642** / **0.327** ±0.242/±0.326 |
| KRAS | 0.664 / **0.299** ±0.259/±0.171 | **0.668** / 0.289 ±0.242/±0.239 | 0.558 / 0.161 ±0.202/±0.085 | 0.589 / 0.188 ±0.198/±0.103 |
| PIK3CA | 0.499 / 0.205 ±0.111/±0.088 | 0.555 / 0.233 ±0.148/±0.079 | **0.601** / **0.323** ±0.162/±0.157 | 0.580 / 0.288 ±0.114/±0.131 |
| RNF43 | 0.588 / 0.279 ±0.188/±0.142 | **0.614** / **0.336** ±0.191/±0.209 | 0.547 / 0.224 ±0.140/±0.124 | 0.554 / 0.247 ±0.149/±0.145 |
| SMAD4 | **0.667** / **0.300** ±0.222/±0.211 | 0.644 / 0.284 ±0.199/±0.166 | 0.505 / 0.160 ±0.250/±0.085 | 0.461 / 0.184 ±0.211/±0.175 |
| TP53 | 0.344 / 0.367 ±0.096/±0.047 | 0.434 / 0.394 ±0.088/±0.045 | 0.581 / 0.515 ±0.100/±0.079 | **0.633** / **0.575** ±0.114/±0.104 |
| **TGCT** (AUROC / AUPRC) | | | | |

Continued on next page

**Supplementary Table 1**: Mutation prediction performance across filtered cancer–gene tasks (continued)

| Task | GigaPath | HistoMethyl GigaPath | Feather | HistoMethyl Feather |
|---|---|---|---|---|
| KIT | 0.531 / 0.313 ±0.236/±0.131 | <u>0.778</u> / <u>0.571</u> ±0.112/±0.208 | 0.646 / 0.352 ±0.152/±0.106 | **0.833** / **0.626** ±0.135/±0.245 |
| KRAS | 0.211 / 0.107 ±0.157/±0.023 | <u>0.667</u> / 0.243 ±0.142/±0.099 | **0.839** / **0.596** ±0.194/±0.364 | 0.666 / <u>0.417</u> ±0.295/±0.377 |
| **CPTAC GBM** (AUROC / AUPRC) | | | | |
| ATRX | 0.485 / 0.312 ±0.312/±0.296 | **0.583** / **0.384** ±0.290/±0.330 | 0.428 / 0.183 ±0.168/±0.056 | <u>0.540</u> / <u>0.373</u> ±0.295/±0.333 |
| EGFR | 0.438 / 0.257 ±0.224/±0.181 | 0.480 / <u>0.308</u> ±0.284/±0.213 | <u>0.492</u> / 0.258 ±0.194/±0.150 | **0.514** / **0.319** ±0.226/±0.197 |
| MUC16 | 0.385 / 0.249 ±0.357/±0.323 | <u>0.689</u> / 0.336 ±0.238/±0.268 | 0.574 / <u>0.349</u> ±0.352/±0.350 | **0.707** / **0.368** ±0.245/±0.303 |
| NF1 | 0.447 / **0.290** ±0.282/±0.263 | <u>0.486</u> / 0.264 ±0.199/±0.139 | 0.483 / 0.272 ±0.262/±0.188 | **0.512** / <u>0.284</u> ±0.261/±0.209 |
| PIK3CA | **0.769** / **0.640** ±0.174/±0.258 | 0.656 / 0.471 ±0.225/±0.253 | 0.426 / 0.302 ±0.236/±0.172 | <u>0.734</u> / <u>0.495</u> ±0.104/±0.147 |
| PTEN | 0.375 / 0.310 ±0.183/±0.112 | 0.398 / 0.318 ±0.196/±0.125 | **0.518** / **0.427** ±0.220/±0.179 | <u>0.436</u> / <u>0.366</u> ±0.180/±0.103 |
| TP53 | **0.824** / **0.728** ±0.130/±0.185 | 0.695 / 0.581 ±0.159/±0.190 | 0.684 / 0.632 ±0.182/±0.189 | <u>0.741</u> / <u>0.672</u> ±0.158/±0.192 |

**Supplementary Table 2** Morphology prediction performance across eligible cohorts. Cells report AUROC / AUPRC, with cross-validation standard deviations shown below each metric. Bold values indicate the best model for each metric within a cohort, and underlined values indicate the second-best model.

| Task | GigaPath | HistoMethyl GigaPath | Feather | HistoMethyl Feather |
|---|---|---|---|---|
| **Morphology prediction** (AUROC / AUPRC) | | | | |
| BRCA | <u>0.886</u> / 0.660<br>±0.055/±0.124 | 0.865 / 0.604<br>±0.051/±0.139 | 0.884 / **0.675**<br>±0.042/±0.109 | **0.907** / <u>0.662</u><br>±0.020/±0.100 |
| COAD | **0.669** / <u>0.321</u><br>±0.068/±0.106 | <u>0.669</u> / 0.290<br>±0.103/±0.113 | 0.658 / **0.364**<br>±0.163/±0.149 | 0.571 / 0.256<br>±0.090/±0.095 |
| KIRC | **0.987** / **0.771**<br>±0.014/±0.216 | <u>0.983</u> / <u>0.694</u><br>±0.020/±0.291 | 0.794 / 0.246<br>±0.148/±0.177 | 0.886 / 0.158<br>±0.045/±0.058 |
| LGG | **0.596** / <u>0.567</u><br>±0.083/±0.059 | 0.567 / 0.558<br>±0.086/±0.052 | <u>0.576</u> / **0.570**<br>±0.090/±0.086 | 0.516 / 0.522<br>±0.080/±0.082 |
| LUAD | 0.656 / 0.472<br>±0.047/±0.080 | 0.668 / 0.460<br>±0.072/±0.110 | <u>0.676</u> / **0.537**<br>±0.070/±0.103 | **0.688** / <u>0.526</u><br>±0.081/±0.093 |
| READ | 0.612 / 0.447<br>±0.175/±0.256 | 0.735 / <u>0.486</u><br>±0.139/±0.155 | **0.756** / **0.486**<br>±0.180/±0.290 | <u>0.750</u> / 0.477<br>±0.157/±0.191 |
| SR386-READ | 0.505 / 0.149<br>±0.176/±0.069 | **0.528** / 0.147<br>±0.198/±0.092 | <u>0.506</u> / **0.246**<br>±0.231/±0.203 | 0.493 / <u>0.224</u><br>±0.167/±0.186 |
| TGCT | <u>0.907</u> / <u>0.933</u><br>±0.194/±0.138 | 0.898 / 0.922<br>±0.193/±0.139 | <u>0.907</u> / <u>0.933</u><br>±0.194/±0.138 | **0.937** / **0.944**<br>±0.144/±0.116 |

**Supplementary Table 3** Overall survival prediction performance across filtered cohorts. Cells report concordance index, with cross-validation standard deviation shown below each value. Bold values indicate the best model within each cohort, and underlined values indicate the second-best model.

| Task | GigaPath | HistoMethyl GigaPath | Feather | HistoMethyl Feather |
|---|---|---|---|---|
| **Overall survival prediction** (C-index) | | | | |
| BRCA | **0.647** ±0.072 | <u>0.621</u> ±0.067 | 0.474 ±0.095 | 0.537 ±0.043 |
| CPTAC GBM | **0.484** ±0.117 | 0.429 ±0.122 | <u>0.472</u> ±0.117 | 0.423 ±0.074 |
| GBM | 0.504 ±0.030 | **0.525** ±0.034 | 0.462 ±0.072 | <u>0.513</u> ±0.023 |
| HNSC | <u>0.544</u> ±0.058 | 0.535 ±0.049 | 0.518 ±0.071 | **0.549** ±0.044 |
| KIRC | **0.651** ±0.061 | <u>0.642</u> ±0.064 | 0.551 ±0.042 | 0.575 ±0.061 |
| KIRP | **0.718** ±0.149 | <u>0.706</u> ±0.098 | 0.647 ±0.159 | 0.555 ±0.126 |
| LGG | **0.572** ±0.057 | 0.508 ±0.082 | <u>0.569</u> ±0.079 | 0.557 ±0.038 |
| LUAD | 0.451 ±0.086 | <u>0.474</u> ±0.067 | **0.500** ±0.062 | 0.458 ±0.065 |
| READ | 0.416 ±0.281 | <u>0.525</u> ±0.347 | 0.373 ±0.307 | **0.702** ±0.268 |
| STAD | <u>0.501</u> ±0.124 | 0.475 ±0.128 | **0.538** ±0.100 | 0.489 ±0.097 |
| TGCT | **0.500** ±0.000 | **0.500** ±0.000 | **0.500** ±0.000 | **0.500** ±0.000 |

**Supplementary Table 4**: DNA methylation beta-value prediction performance. Each cancer-specific block reports hybrid-site DNA methylation prediction metrics for the four slide encoders. Values are means with cross-validation standard deviations below. Bold values indicate the best model for each metric, and underlined values indicate the second-best model; lower values are preferred for error metrics.

| Metric | GigaPath | HistoMethyl GigaPath | Feather | HistoMethyl Feather |
|---|---|---|---|---|
| **BRCA** (mean / standard deviation) | | | | |
| Flat PCC | <u>0.549</u> ±0.014 | 0.545 ±0.012 | 0.508 ±0.016 | **0.550** ±0.017 |
| Mean site PCC | <u>0.311</u> ±0.018 | 0.291 ±0.015 | 0.270 ±0.010 | **0.323** ±0.015 |
| Median site PCC | <u>0.310</u> ±0.020 | 0.293 ±0.015 | 0.268 ±0.012 | **0.323** ±0.017 |
| Sites PCC > 0.4 | <u>0.296</u> ±0.042 | 0.253 ±0.039 | 0.205 ±0.024 | **0.311** ±0.038 |
| MAE | <u>0.144</u> ±0.002 | 0.144 ±0.002 | 0.150 ±0.003 | **0.143** ±0.003 |
| RMSE | **0.183** ±0.002 | <u>0.183</u> ±0.002 | 0.191 ±0.004 | 0.183 ±0.004 |
| **KIRC** (mean / standard deviation) | | | | |
| Flat PCC | <u>0.630</u> ±0.021 | 0.624 ±0.026 | 0.583 ±0.021 | **0.635** ±0.025 |
| Mean site PCC | <u>0.404</u> ±0.012 | 0.378 ±0.018 | 0.355 ±0.023 | **0.427** ±0.017 |
| Median site PCC | <u>0.406</u> ±0.016 | 0.382 ±0.021 | 0.348 ±0.025 | **0.429** ±0.018 |
| Sites PCC > 0.4 | <u>0.510</u> ±0.022 | 0.475 ±0.034 | 0.419 ±0.035 | **0.547** ±0.027 |
| MAE | <u>0.106</u> ±0.005 | 0.106 ±0.006 | 0.113 ±0.006 | **0.105** ±0.006 |
| RMSE | <u>0.141</u> ±0.007 | 0.141 ±0.008 | 0.150 ±0.007 | **0.140** ±0.007 |
| **OV** (mean / standard deviation) | | | | |
| Flat PCC | 0.434 ±0.009 | **0.450** ±0.012 | 0.388 ±0.032 | <u>0.448</u> ±0.008 |
| Mean site PCC | 0.089 ±0.015 | 0.074 ±0.020 | <u>0.092</u> ±0.012 | **0.153** ±0.010 |
| Median site PCC | 0.088 ±0.016 | 0.074 ±0.021 | <u>0.090</u> ±0.013 | **0.152** ±0.009 |
| Sites PCC > 0.4 | 0.004 ±0.002 | <u>0.005</u> ±0.003 | 0.004 ±0.002 | **0.022** ±0.005 |
| MAE | 0.195 ±0.003 | <u>0.193</u> ±0.003 | 0.200 ±0.005 | **0.192** ±0.003 |
| RMSE | 0.235 ±0.003 | **0.233** ±0.003 | 0.247 ±0.012 | <u>0.235</u> ±0.003 |

Continued on next page

**Supplementary Table 4**: DNA methylation beta-value prediction performance (continued)

| Metric | GigaPath | HistoMethyl GigaPath | Feather | HistoMethyl Feather |
|---|---|---|---|---|
| **UCEC** (mean / standard deviation) | | | | |
| Flat PCC | 0.528 ±0.029 | 0.501 ±0.024 | 0.499 ±0.032 | **0.538** ±0.020 |
| Mean site PCC | 0.276 ±0.035 | 0.211 ±0.013 | 0.265 ±0.034 | **0.317** ±0.020 |
| Median site PCC | 0.275 ±0.040 | 0.218 ±0.021 | 0.261 ±0.035 | **0.317** ±0.020 |
| Sites PCC > 0.4 | 0.241 ±0.078 | 0.152 ±0.037 | 0.188 ±0.068 | **0.295** ±0.052 |
| MAE | 0.160 ±0.005 | 0.165 ±0.003 | 0.165 ±0.005 | **0.158** ±0.003 |
| RMSE | 0.200 ±0.005 | 0.203 ±0.003 | 0.207 ±0.005 | **0.199** ±0.003 |
| **LUAD** (mean / standard deviation) | | | | |
| Flat PCC | 0.524 ±0.023 | 0.512 ±0.020 | 0.512 ±0.037 | **0.552** ±0.026 |
| Mean site PCC | 0.254 ±0.021 | 0.185 ±0.027 | 0.282 ±0.035 | **0.328** ±0.029 |
| Median site PCC | 0.257 ±0.021 | 0.191 ±0.024 | 0.289 ±0.037 | **0.336** ±0.033 |
| Sites PCC > 0.4 | 0.210 ±0.052 | 0.144 ±0.044 | 0.243 ±0.090 | **0.356** ±0.077 |
| MAE | 0.124 ±0.004 | 0.125 ±0.003 | 0.127 ±0.004 | **0.120** ±0.004 |
| RMSE | 0.157 ±0.005 | 0.157 ±0.004 | 0.163 ±0.007 | **0.154** ±0.005 |
| **THCA** (mean / standard deviation) | | | | |
| Flat PCC | 0.481 ±0.004 | 0.462 ±0.005 | 0.458 ±0.004 | **0.507** ±0.007 |
| Mean site PCC | 0.171 ±0.005 | 0.136 ±0.010 | 0.184 ±0.006 | **0.235** ±0.013 |
| Median site PCC | 0.126 ±0.006 | 0.103 ±0.010 | 0.147 ±0.010 | **0.196** ±0.016 |
| Sites PCC > 0.4 | 0.137 ±0.006 | 0.110 ±0.010 | 0.138 ±0.007 | **0.197** ±0.015 |
| MAE | 0.143 ±0.001 | 0.145 ±0.001 | 0.147 ±0.001 | **0.139** ±0.002 |
| RMSE | 0.180 ±0.001 | 0.181 ±0.001 | 0.186 ±0.001 | **0.177** ±0.002 |
| **GBM** (mean / standard deviation) | | | | |
| Flat PCC | 0.412 ±0.015 | **0.449** ±0.012 | 0.377 ±0.041 | 0.425 ±0.016 |

Continued on next page

**Supplementary Table 4**: DNA methylation beta-value prediction performance (continued)

| Metric | GigaPath | HistoMethyl GigaPath | Feather | HistoMethyl Feather |
|---|---|---|---|---|
| Mean site PCC | 0.041 ±0.020 | 0.031 ±0.013 | <u>0.091</u> ±0.055 | **0.112** ±0.017 |
| Median site PCC | 0.041 ±0.020 | 0.030 ±0.014 | <u>0.090</u> ±0.055 | **0.109** ±0.016 |
| Sites PCC > 0.4 | 0.002 ±0.002 | 0.002 ±0.001 | <u>0.010</u> ±0.008 | **0.012** ±0.005 |
| MAE | 0.184 ±0.002 | **0.180** ±0.002 | 0.194 ±0.005 | <u>0.183</u> ±0.001 |
| RMSE | 0.226 ±0.002 | **0.219** ±0.003 | 0.240 ±0.005 | <u>0.226</u> ±0.001 |
| **LGG** (mean / standard deviation) | | | | |
| Flat PCC | 0.486 ±0.018 | <u>0.496</u> ±0.005 | 0.472 ±0.033 | **0.508** ±0.019 |
| Mean site PCC | 0.224 ±0.018 | 0.215 ±0.034 | <u>0.228</u> ±0.036 | **0.270** ±0.022 |
| Median site PCC | 0.236 ±0.022 | <u>0.236</u> ±0.040 | 0.235 ±0.037 | **0.279** ±0.022 |
| Sites PCC > 0.4 | <u>0.198</u> ±0.049 | 0.192 ±0.091 | 0.169 ±0.074 | **0.243** ±0.062 |
| MAE | 0.171 ±0.005 | <u>0.170</u> ±0.004 | 0.175 ±0.007 | **0.169** ±0.004 |
| RMSE | 0.213 ±0.005 | **0.209** ±0.004 | 0.220 ±0.008 | <u>0.211</u> ±0.005 |

### 2 Pre-training

#### 2.1 Pre-training data

Contrastive pre-training used 9,607 paired whole-slide image–DNA methylation records from 8,508 TCGA [1] cases across 29 TCGA projects. Each record contained a tiled H&E slide directory and a matched DNA methylation beta-value file for the same case. No downstream labels were used during this stage. LGG was absent from the pre-training index and was therefore reserved for disease-held-out evaluation.

For each methylation profile, CpG beta-value files were parsed as CpG-site identifiers and continuous beta values. Missing, non-finite, or non-numeric beta values were removed. CpG sites were then intersected with the CpGPT [2] large vocabulary, which contained 1,187,190 CpG sites in the reported setup. The methylation branch used CpGPT-large as a frozen encoder, followed by a trainable MLP projection head.

#### 2.2 Image and methylation encoders

Two HistoMethyl variants were trained, one initialized from Prov-GigaPath [3] and one initialized from Feather-24k [4]. In both variants, tile-level image encoders were frozen. The trainable image-side components were the slide encoder and the image projection head. The image branch first aggregated tile features and tile coordinates into a slide-level embedding, then projected the slide embedding into the contrastive embedding space using an MLP head.

The methylation branch used frozen CpGPT-large embeddings followed by a trainable methylation projection head. Both the image and methylation projection heads used an MLP structure with hidden dimension equal to twice the output embedding dimension, dropout 0.2, and layer normalization. The GigaPath-based model used a 768-dimensional shared embedding space; the Feather-based model used a 256-dimensional shared embedding space.

#### 2.3 Contrastive objective and optimization

Let $u_i$ denote the projected image embedding and $v_i$ denote the projected methylation embedding for matched case $i$. Pairwise similarity was computed as

$$s_{ij} = \frac{u_i^\top v_j}{\tau},$$

where $\tau$ was initialized to 0.07 through a learnable logit scale. The GigaPath-based model and Feather-based model were trained with a SigLIP-style sigmoid contrastive loss. In both cases, the objective increased similarity for matched slide–methylation pairs and decreased similarity for unmatched pairs in the minibatch. The logit scale was clamped after backpropagation to keep the effective scale within a bounded range.

#### 2.4 Representation visualization

For the t-SNE visualization, the same set of slides was used across all four encoders: GigaPath, HistoMethyl GigaPath, Feather, and HistoMethyl Feather. The visualization included the 12 displayed cancer groups, with at most 80 slides sampled per cancer group using random seed 17. Slide embeddings were standardized, reduced by PCA to at most 50 dimensions, and then embedded into two dimensions with t-SNE using Euclidean distance, PCA initialization, perplexity 35, automatic learning rate, random seed 17, and 1,500 iterations. The resulting two-dimensional coordinates were standardized before plotting. The t-SNE analysis was used only to visualize representation structure and was not used for downstream training, model selection, or cohort selection.

#### 2.5 Image–methylation retrieval

Cross-modal retrieval was used to evaluate whether the learned embedding space aligned H&E slides with matched methylation profiles before supervised downstream adaptation. Retrieval was evaluated on GBM, HNSC, and BRCA samples from the reconstructed pre-training validation split and on LGG samples from the held-out evaluation set. Each cohort contributed 38 paired slide-methylation samples, to ensure consistent pool size, selected using random seed 42. For each cohort, the task was to retrieve the matched DNAm profile for each slide from the cohort-specific methylation pool.

**Supplementary Table 5** Pre-training hyperparameters for the two HistoMethyl variants.

| Setting | HistoMethyl GigaPath | HistoMethyl Feather |
|---|---|---|
| Image backbone | Prov-GigaPath | Feather-24k |
| Methylation backbone | CpGPT-large | CpGPT-large |
| Tile encoder | Frozen | Frozen |
| Slide encoder | Trainable | Trainable |
| CpGPT encoder | Frozen | Frozen |
| Projection head | MLP | MLP |
| Embedding dimension | 768 | 256 |
| Initial temperature | 0.07 | 0.07 |
| Optimizer | AdamW | AdamW |
| Learning rate | $1 \times 10^{-4}$ | $1 \times 10^{-4}$ |
| Weight decay | 0.01 | 0.01 |
| AdamW betas | 0.9, 0.95 | 0.9, 0.95 |
| Batch size | 48 | 128 |
| Maximum epochs | 30 | 100 |
| Warmup steps | 780 | 780 |
| Gradient clipping | 5 | 5 |
| Precision | bfloat16 mixed | bfloat16 mixed |
| Validation fraction | 10% | 10% |
| Pre-training seed | 24442 | 24442 |

For HistoMethyl models, the slide embedding was the methylation-aligned image embedding. For image-only baselines, the original slide embedding was passed through the corresponding trained image projection head so that baseline and HistoMethyl embeddings were compared in the same projected space. CpGPT methylation embeddings were passed through the trained methylation projection head. Image and methylation embeddings were L2-normalized, and retrieval was performed by cosine similarity. Recall@1, Recall@5, and Recall@10 were computed as the fraction of slides whose matched methylation profile appeared within the top $k$ retrieved methylation profiles. The main figure reports image-to-DNAm retrieval.

### 3 Downstream Task Adaptation

#### 3.1 Cross-validation and aggregation

For mutation prediction, receptor-associated prediction, morphology-associated classification, and survival prediction, downstream evaluation used cohort-specific five-fold cross-validation repeated with random seeds 42, 43, and 44. Thus, each reported classification or survival task was summarized over 15 held-out folds when all folds were valid. For classification tasks, folds were stratified by the target label. When more than one slide was available from the same case, splitting was grouped by case identifier so that slides from the same case could not appear in both training and test folds. Survival folds were also grouped by case identifier when applicable, but were not label-stratified because the endpoint was right-censored time-to-event data.

All four encoders were evaluated under identical splits and task heads within each endpoint. HistoMethyl GigaPath was compared only against the matched GigaPath baseline, and HistoMethyl Feather was compared only against the matched Feather baseline. For aggregate mutation summaries, AUROCs were first averaged across mutation genes within each cohort and then averaged equally across cohorts, so larger gene panels or larger cohorts did not dominate the reported pan-cancer mean.

#### 3.2 Classification probes

Mutation prediction, BRCA receptor-associated prediction, and morphology-associated prediction were treated as binary slide-level classification tasks. The input to each task was a fixed slide embedding. The classifier was a one-hidden-layer MLP consisting of a linear layer to 256 hidden units, ReLU activation, and a final linear layer with one output logit. The model was trained with binary cross-entropy with logits. Predicted positive-class probabilities were obtained with the sigmoid function.

All classification probes used Adam optimization with learning rate $3 \times 10^{-4}$, weight decay 0, batch size 64, and 50 epochs. The same architecture, optimizer, and training schedule were used for all mutation, receptor, and morphology tasks. AUROC was the primary metric used for the main figures, and AUPRC was retained in the supplementary tables.

#### 3.3 Survival probes

Overall survival prediction used fixed slide embeddings as input to a linear Cox proportional hazards model. Survival time was taken from overall survival days when available; otherwise, days to death and days to last follow-up were used as fallback time fields. The event indicator was derived from the survival event annotation, with vital status used to identify deaths when available. Samples without usable time or event information were excluded from that survival endpoint.

Before Cox fitting, slide embeddings were standardized using the training fold. The Cox model used a linear risk score and was trained by minimizing the negative Cox partial log-likelihood. Optimization used Adam with learning rate $1 \times 10^{-2}$, weight decay $1 \times 10^{-4}$, and 200 epochs. The trained model produced one risk score per held-out slide, where higher scores indicated higher predicted risk. Performance was summarized by Harrell's C-index on the held-out fold and then averaged across folds and seeds.

#### 3.4 DNA methylation beta-value prediction

DNA methylation beta-value prediction was evaluated as multi-output regression from H&E slide representations to continuous CpG beta values. This analysis used five-fold cross-validation with random seed 42. CpG target selection was performed independently inside each training fold and used only training-fold methylation files. Following the DEPLOY methylation prediction strategy for selecting informative beta-value targets and reporting both global and site-wise prediction metrics [5], CpG sites were restricted to CpGPT-vocabulary sites with adequate observation frequency and variability in the training split.

Specifically, a CpG site was eligible if it was observed in at least 95% of training methylation profiles, had training-set beta-value standard deviation of at least 0.08, and was not dominated by a single methylation state. To remove nearly constant binary-like sites, beta values were thresholded at 0.5, and a site was excluded if more than 80% of observed training values fell on the same side of that threshold. All available training profiles were used to compute these statistics. No held-out fold methylation values were used to select CpG sites.

For each fold, selected CpG sites were assembled into a target matrix with one row per slide and one column per CpG site. A binary observation mask was stored in parallel so that missing beta values did not contribute to training or evaluation. The beta-value predictor used the slide encoder followed by a regression head with a 256-dimensional hidden layer, ReLU activation, and one output per selected CpG site. The slide encoder and regression head were trained jointly. HistoMethyl regression models loaded the corresponding contrastively pre-trained slide encoder before training the regression head.

The regression head was optimized with AdamW using learning rate $2 \times 10^{-4}$, weight decay 0, betas 0.9 and 0.95, and batch size 64. GigaPath-based models were trained for 30 epochs, and Feather-based models were trained for 70 epochs. Within each training fold, 5% of training samples were used as an internal validation subset, and the model state with the lowest validation masked mean squared error was retained. The loss was masked mean squared error:

$$\mathcal{L}_{\text{masked}} = \frac{\sum_{i,j} m_{ij}(\hat{y}_{ij} - y_{ij})^2}{\sum_{i,j} m_{ij}},$$

where $y_{ij}$ is the measured beta value for sample $i$ and CpG site $j$, $\hat{y}_{ij}$ is the predicted beta value, and $m_{ij}$ indicates whether that beta value was observed.

Metrics were computed on observed held-out beta values only. Flattened Pearson correlation was computed after flattening all observed sample–CpG pairs in the held-out fold. Site-wise Pearson correlation was computed separately for each CpG site with sufficient observed values and nonzero variance, then summarized by the mean and median across sites. Mean absolute error, mean squared error, and root mean squared error were computed with the same observation mask. The main figure reports flattened Pearson correlation, median site-wise Pearson correlation, and mean absolute error; the supplementary table includes the broader metric set.

#### 3.5 Occlusion-based mutation heatmaps

Occlusion heatmaps were generated for selected mutation prediction examples to visualize which local tissue regions increased the positive-class mutation score. The heatmap analysis used the same four encoders shown in the quantitative experiments. Candidate slides were required to have at least 300 tiled patches and at least 220 retained tissue patches after tissue masking so that the rendered heatmap was sufficiently dense. Slides with long-to-short aspect ratio greater than 2.0 were excluded from automatic selection to avoid visually distorted panels. The number of rendered examples was capped per project and per slide.

For each selected project–gene–slide example, the slide was assigned to a held-out fold using five-fold stratified splitting with seed 42. A lightweight logistic regression mutation probe was trained on the training samples from that fold using fixed slide embeddings from the corresponding encoder. The selected slide was kept in the held-out fold. The original slide embedding was computed from tile features and spatial coordinates, and the mutation probe positive-class decision score was recorded as the baseline score.

Tissue masking was performed at the tile level. A tile was retained if the estimated tissue fraction was at least 0.22. The default segmentation backend used a local HEST tissue segmentation model when available and otherwise used a heuristic based on grayscale intensity and color saturation. In the heuristic path, pixels were treated as tissue-like when mean intensity was below 0.92 or saturation exceeded 0.08. The tile tissue mask was eroded by one tile-neighborhood step to reduce edge and background spillover.

Occlusion was performed over spatial groups of tissue patches rather than single patches. Retained tissue patches were assigned to a regular spatial grid. The grid size was chosen dynamically from the number of patches, targeting approximately two patches per region, with the number of bins constrained between 10 and 28 per axis. For each region, tile features in that region were replaced by the mean tile feature vector from the same slide. The slide embedding was then recomputed with the same slide encoder, and the mutation probe decision score was recomputed. The regional importance score was defined as the decrease in positive-class decision score after masking:

$$\Delta_r = f(x) - f(x_{\setminus r}),$$

where $f(x)$ is the baseline positive-class decision score and $f(x_{\setminus r})$ is the score after replacing region $r$ with mean tile features. Negative decreases were set to zero, so only regions whose removal reduced the positive mutation score contributed to the overlay.

For rendering, regional scores were assigned back to patch locations. Scores were normalized within each displayed slide and model using the 97th percentile of nonzero tissue-region values, then gamma-adjusted with $\gamma = 0.85$. Because normalization was performed within each panel, these heatmaps are intended for qualitative comparison of spatial evidence patterns rather than quantitative comparison of absolute attribution magnitude across slides.

### References

[1] Tomczak, K., Czerwińska, P., Wiznerowicz, M.: Review the cancer genome atlas (tcga): an immeasurable source of knowledge. Contemporary Oncology/Współczesna Onkologia **2015**(1), 68–77 (2015)

[2] Lima Camillo, L.P., Sehgal, R., Armstrong, J., Miller, H.E., Lasky-Su, J.A., Higgins-Chen, A.T., Horvath, S., Wang, B.: Cpgpt: a foundation model for dna methylation. bioRxiv, 2024–10 (2024)

[3] Xu, H., Usuyama, N., Bagga, J., Zhang, S., Rao, R., Naumann, T., Wong, C., Gero, Z., González, J., Gu, Y., Xu, Y., Wei, M., Wang, W., Ma, S., Wei, F., Yang, J., Li, C., Gao, J., Rosemon, J., Bower, T., Lee, S., Weerasinghe, R., Wright, B.J., Robicsek, A., Piening, B., Bifulco, C., Wang, S., Poon, H.: A whole-slide foundation model for digital pathology from real-world data. Nature (2024)

[4] Shao, D., Chen, R.J., Song, A.H., Runevic, J., Lu, M.Y., Ding, T., , Mahmood, F.: Do multiple instance learning models transfer? In: International Conference on Machine Learning (2025)

[5] Hoang, D.-T., Shulman, E.D., Turakulov, R., Abdullaev, Z., Singh, O., Campagnolo, E.M., Lalchungnunga, H., Stone, E.A., Nasrallah, M.P., Ruppin, E., *et al.*: Prediction of dna methylation-based tumor types from histopathology in central nervous system tumors with deep learning. Nature Medicine **30**(7), 1952–1961 (2024)
